# Red Blood Cell-derived Extracellular Vesicles enable Cisplatin and Cetuximab combined Therapy against Triple-Negative Breast Cancer

**DOI:** 10.1101/2025.03.20.644320

**Authors:** Miriam Romano, Angelo Musicò, Andrea Zendrini, Enrica Gilberti, Francesca Orlandi, Tatiana Pedrazzi, Silvia Alacqua, Agnese Segala, Rossella Zenatelli, Selene Tassoni, Lucia Paolini, Ingrid Cifola, Eleonora Mangano, Clarissa Consolandi, Tania Camboni, Lorena Signati, Serena Mazzucchelli, Arabella Neva, Maurizio Ragni, Marco Severgnini, Alessandra Valerio, Giuseppe Pomarico, Camillo Almici, Giuseppe De Palma, Fabio Corsi, Paolo Bergese, Annalisa Radeghieri

## Abstract

**Background:** Triple-negative breast cancer is an aggressive breast cancer subtype characterized by the absence of human epidermal growth factor receptor 2, estrogen and progesterone receptors, limiting targeted therapy options. Cisplatin, a chemotherapeutic agent, induces DNA damage and exhibits some efficacy against triple-negative breast cancer, but its effectiveness is often reduced by chemoresistance and systemic toxicity.

A very promising strategy to augment cisplatin treatment can be based on combining it with the biologic Cetuximab, an epidermal growth factor receptor inhibitor, which boosts cisplatin efficacy by inducing ferroptosis.

**Results:** To optimize this strategy in a biocompatible and precise manner, we developed a nanoplatform based on red blood cell-derived extracellular vesicles for the combined delivery of Cetuximab and cisplatin, enabling immune evasion, and the possibility of autologous personalization and GMP-compliant production. Owing to their DNA-free lumen and lack of EGFR, RBC-EVs preserve cisplatin activity and prevent interference with cetuximab. This formulation enhances cisplatin’s cytotoxicity by up to 50%, as shown *in vitro* and in patient-derived organoids. It effectively reduces chemoresistance by downregulating hypoxia-related genes and promoting ferroptosis, additionally, it improves cisplatin’s cytotoxic effects while reducing hemotoxicity compared to the administration of free cisplatin.

**Conclusions:** These findings highlight the potential of red blood cell-derived extracellular vesicles as a biocompatible delivery system enabling combined therapy and offering a promising strategy to overcome current limitations in TNBC treatment.

## Background

Triple-negative breast cancer (TNBC) is one of the most aggressive breast cancer subtypes, generally characterized by the lack of expression of human epidermal growth factor receptor 2 (HER2), estrogen receptors (ER), and progesterone receptors (PR), making it resistant to many targeted therapies (1). Recent studies highlighted TNBC sensitivity to platinum-based drugs (e.g. carboplatin, cisplatin), particularly due to defects in DNA repair mechanisms, such as those mediated by the homologous recombination (HR) pathway (2). Such drugs, by forming intra- and inter-crosslinks with the DNA, can induce DNA damage, leading to apoptosis and, ultimately, cell death. Among platinum-based drugs, cisplatin is receiving increased interest for TNBC treatment, alone or in combination with other physical (e.g. surgical intervention (3)) or chemical (e.g. gemcitabine or nab-paclitaxel (4)) treatment options. Cisplatin is a small hydrophilic drug with remarkable cytotoxic activity and exerts its anti-cancer effects primarily in the nuclei (2). However, cisplatin efficacy in TNBC is weakened by the activation of various cellular resistance mechanisms, including alterations of drug influx-efflux routes and detoxification processes (5), that might hamper its nuclear activity. Recent findings reported that ferroptosis, a form of regulated cell death triggered by an iron-dependent accumulation of reactive oxygen species (ROS) and lipid peroxidation products, can increase the sensitivity of TNBC cells to cisplatin (6–8). In fact, Cisplatin can interact with reduced glutathione (GSH), impairing the activity of the glutathione peroxidase 4 (GPX4) in neutralizing harmful lipid peroxides. This disruption shifts the cell redox balance to oxidative stress and cell death by ferroptosis (6,9). However, the “hypoxia status” and therefore ferroptosis can be inhibited through the stabilization of the hypoxia-inducible factor 1-alpha (HIF-1α) transcription factor (10,11). In this context, TNBC frequently over-expresses the epidermal growth factor receptor (EGFR), which further activates HIF-1α. This activation helps TNBC cells to survive in hypoxic environments, thus contributing to cisplatin resistance (6,12,7). Cetuximab (CTX, Erbitux^®^), an FDA-approved anti-EGFR monoclonal antibody (13,14), may represent a promising treatment option in this case, since it has been designed to bind the extracellular domain of EGFR and inhibit its downstream signaling cascade. This consequently also leads to the down-regulation of HIF-1α as already showed for head and neck squamous cell carcinoma (15,16) and TNBC. (17). Therefore, the combined use of cisplatin and CTX may offer several advantages, including an enhanced cisplatin anti-tumor activity, the ability to overcome drug resistance mechanisms and cisplatin action as pro-ferroptosis agent (18).

Currently, combined therapies use cisplatin and CTX separately, leading to overdosing and off-target toxicity. In this scenario, nanotechnology may offer a solution by incorporating both agents into a single nanoplatform, thus enabling their simultaneous delivery and improving selectivity while reducing off-target effects (19). Moreover, the use of biocompatible nano drug delivery systems (DDS) may further help in minimizing the systemic toxicity and limiting the immune system activation. In this context, extracellular vesicles (EVs) are biogenic nanoparticles that have gained more and more interest as innovative delivery systems due to their intrinsic ability to carry biological payloads and to be recognized as self by the body. EVs are composed of a biological membrane enclosing and protecting an aqueous core containing proteins, nucleic acids and metabolites (20). Several types of EVs have already been proposed as DDS for cancer therapy, and in particular cisplatin delivery by EVs has already been attempted with milk-derived EVs against resistant ovarian cancer (21). Among the plethora of EV cell types, red blood cell-derived EVs (RBC-EVs) are a highly promising next-generation drug delivery nanoplatform, offering an optimal balance of physicochemical and biological properties, superior scalability, and clinical translation potential over lipid nanoparticles (LNPs) and other EVs (22,23). They boast high-yield, reproducible production under established GMP standards, biocompatibility, immune evasion, and minimal oncogenic risk (23,24). Their ability to be derived from a patient’s own RBCs enables a truly personalized, autologous nanomedicine approach. Beyond their favorable properties, RBC-EVs may offer a mechanistic advantage for cisplatin delivery. Their DNA-free lumen should minimize non-productive platinum-nucleophile interactions, preserving the drug in a functionally competent state during circulation and favoring activation only upon uptake by tumor cells (25,26).

Additionally, RBC-EVs lack EGFR, thus preventing interference with CTX during surface engineering (27). These properties make RBC-EVs the ideal basis for the development of a multimodal nanoplatform for drug delivery.

In this study, we developed a nanoformulation based on human RBC-EVs loaded with cisplatin and surface-functionalized with CTX (^CTX^RBC-EV_cispt_). ^CTX^RBC-EV_cispt_ demonstrated the ability to enhance cisplatin uptake *in vitro* and in patient-derived organoids (PDOs). It increased cytotoxicity and reduced chemoresistance by down-regulating hypoxia-related genes, offering superior outcomes as compared to the administration of free cisplatin. Additionally, ^CTX^RBC-EV_cispt_ mitigated *ex vivo* toxicity towards red blood cells.

## Methods

### Production and separation of RBC-EVs

We have submitted all relevant data from our experiments to the EV-TRACK knowledge base (EV-TRACK EV250024) (28). Red blood cells (RBCs) collected from anonymous Type 0+ healthy donors, who provided written consent, were supplied by the Blood Transfusion Unit of Spedali Civili Hospital of Brescia (ethical approval NP5705) in sealed, sterile bags. RBC-EVs were produced by chemical induction of RBC with CaCl_2_ and calcium ionophore, as previously described (27,29). Briefly, 300 mL of RBC concentrates were centrifuged (5804R Eppendorf centrifuge equipped with an A-4-44 rotor, Eppendorf, Hamburg, Germany) at 1000x*g* for 8 minutes at 4°C and washed twice in PBS without CaCl_2_ and MgCl_2_. The RBC pellets were then resuspended with PBS without CaCl_2_ and MgCl_2_ but supplemented with 0.1 g/L of CaCl_2_ (CPBS, ratio 2 CPBS : 1 RBCs) and centrifuged at 1000x*g* for 8 minutes at 4°C. RBCs in CPBS were transferred into 175 mm^2^ tissue culture flasks (Sarstedt Ag & Co. KG, Nümbrecht, Germany) and incubated at 37°C (5% CO_2_) for 15 h with calcium ionophore (final concentration 10 µM). RBCs were diluted 1:1 with PBS without CaCl_2_ and MgCl_2_ and RBC-EVs were obtained by multiple sequential centrifugation steps, all carried out at 4°C (600x*g* for 20 minutes, 1600x*g* for 15 minutes, 3200x*g* for 15 minutes, with the 5804R Eppendorf centrifuge equipped with an A-4-44 rotor, Eppendorf, Hamburg, Germany, and 10000x*g* for 30 minutes with the Optima XPN-100 equipped with a TY45 Ti rotor, Beckman Coulter, Brea, CA, USA). In each step, the supernatant was processed, while the pellets were discarded. The 10000x*g* supernatants were filtered (Primo syringe 0.45 µm filter, EPVPE45250, Euroclone S.p.A., Milan, Italy) and ultracentrifuged at 50,000x*g* for 70 minutes at 4°C (Optima XPN-100 equipped with a TY45 Ti rotor, Beckman Coulter, USA). 2 mL of RBC-EVs were layered above 2 mL of cold PBS without CaCl_2_ and MgCl_2_ added on top of frozen 60% sucrose (1 mL) and ultracentrifuged at 50000x*g* for 16 h at 4°C with deceleration speed set to 0 (Optima MAX-XP equipped with a MLS-50 rotor, Beckman Coulter, USA). The red layers of RBC-EVs were collected, resuspended in PBS without CaCl_2_ and MgCl_2_ and ultracentrifuged at 50000x*g* for 70 minutes at 4°C (Optima MAX-XP equipped with a TLA-55 rotor, Beckman Coulter, USA). Finally, the RBC-EV pellets were resuspended in 1 mL of cold PBS without CaCl_2_ and MgCl_2_ and stored at – 80°C until use.

### Biophysical and biochemical characterization of RBC-EVs

The purity of RBC-EVs from soluble contaminants and cellular residues was checked with the CONAN assay, following the protocol previously described (30). Nanoparticle tracking analysis (NTA) was performed according to the manufacturer’s instructions using a NanoSight NS300 system configured with a 532 nm laser (Malvern Panalytical, Malvern, UK). The samples were diluted 1:1000 in filtered PBS (0.22 μm) to a final volume of 1 mL to obtain the optimal particle per frame value (20–100 particles/frame). A syringe pump with constant flow injection was used (20 μL·min−1), and the temperature was set constant at 25 °C. Particles were detected at a camera level of 10 and three videos of 60 s were captured and analyzed with NTA software (version 3.2, Malvern Panalytical). Atomic force microscopy (AFM) imaging was performed on a Nanosurf NaioAFM equipped with a Multi75-AI-G tip (Budget Sensors, Sofia, Bulgaria). For sample preparation, RBC-EVs were resuspended in 100 μL sterile H_2_O (Milli-Q, Merck Millipore, Burlington, MA, USA) and diluted 1:10 in H_2_O. 5 μL of the samples were then spotted onto freshly cleaved mica sheets (Grade V-1, thickness 0.15 mm, size 15 × 15 mm2) and dried at 50 °C for 10 minutes. Images were acquired in tapping mode, with a scan size ranging from 1.5 to 25 μm and a scan speed of 1 s per scanning line. Image processing was performed on Gwyddion software (version 2.61). Protein concentrations of RBC-EVs and RBC homogenate samples were determined with a Pierce™ BCA Protein Assay Kit (ThermoFisher, Rockford, IL, USA), following the manufacturer’s instructions. RBC-EV biochemical characterization was carried out with SDS-PAGE followed by Western Blot analysis. Samples were mixed with 6X Laemmli buffer and boiled for 5 min at 95°C. 30µg of proteins were separated by SDS-PAGE (10% polyacrylamide) and transferred onto a PVDF membrane. The blocking step was carried out with a 5% fat-free milk in PBS 0.05% Tween-20 (PBS-T) for 1h at 37°C. Membranes were incubated overnight at 4°C with the following primary antibodies diluted in 1% fat-free milk PBS-T: anti-BAND3 (1:1000, clone A-6, sc-133190, Santa Cruz Biotechnology, Dallas, TX, USA), anti-Flotillin (1:500, clone C-2, sc-74566, Santa Cruz Biotechnology, USA) and anti-Hemoglobin subunit β (HBB, 1:500, H00003043-M02A, Abnova, Taiwan). The membranes were washed thrice for 10 min with PBS-T and incubated for 1 h with rabbit anti-mouse HRP conjugated secondary antibody diluted 1:3000 in 1% fat-free milk PBS-T (Bethyl Laboratories, Montgomery, TX, USA). Images were acquired with Chemibox (Syngene, Cambridge, UK).

### Membrane engineering of RBC-EVs with Cetuximab

Cetuximab antibody was conjugated onto RBC-EVs and RBC-EV_cispt_ membrane via the strain-promoted alkyne–azide cycloaddition (SPAAC) reaction, as previously described. Briefly, CTX storage buffer was replaced with 0.1 mM NaHCO_3_, pH 8.4, using VivaSpin2000 column with 50 kDa cut-off. CTX solutions were centrifuged at 2000 xg for 15 minutes. The buffer exchange step was carried out thrice by adding 2 mL of NaHCO_3_ 0.1 mM, pH 8.4 at each step. CTX concentration after the buffer exchange process was quantified by NanoDrop™ OneC (Thermo Fisher Scientific, Waltham, MA, USA). For antibody functionalization, 200 µL of CTX 0.016 mM was incubated overnight on continuous mixing at 4 °C with 12 equivalents of Sulfo Cyanine 7.5 NHS ester (0.2 mM in DMSO) and 6 equivalents of DBCO STF ester (0.1 mM in DMSO), thus obtaining modified CTX. RBC-EV samples were reacted with 2000 equivalents of PEG4-NHS-ester (diluted in PBS) overnight with continuous mixing at 4 °C. PEG excess was removed using a Vivaspin 500 column with a 10 kDa cut-off, washing the samples five times with 500 μL of PBS. Pegylated RBC-EVs were recovered from the column with 200 μL of PBS. 200 μL of pegylated RBC-EVs (8 × 10^11^ particles per mL) were then reacted overnight with continuous mixing at 4 °C with 2000 equivalents of modified CTX (in PBS, pH 7.4). The unreacted antibody was removed by ultracentrifugation (100000 xg, 2 h), and the pellet was resuspended in 100 μL of PBS and stored at 4 °C until use. For CTX quantification, known amounts (2µg, 0.6µg, 0.2µg, 0.06µg, 0.02µg) of Cy7.5-CTX were loaded onto a Any kD™ Mini-PROTEAN TGX Stain-Free precast gel for polyacrylamide gel electrophoresis (PAGE) (Bio-Rad, Hercules, CA, USA). After the electrophoresis run, the gel was imaged by G:BOX Chemi XX9 (Syngene), with an acquisition time of 2 min and 800 nm wavelength. The image was analyzed using Image Lab software (version 6.0, Bio-Rad). For the optimization of the RBC-EV membrane engineering, results related to CTX molecules per EVs from eight different preparations were used and expressed as mean ± standard error of the mean (SEM).

### Cisplatin encapsulation and quantification

200 µL of RBC-EVs or ^CTX^RBC-EVs (2 x 10^12^ EVs/mL) were incubated with 200 µL of 1.4 mg/L cisplatin (Sandoz AG, Novartis, Basel, Switzerland) for 4h or 24h at 37°C in agitation on a tubes stirrer. Cisplatin-loaded RBC-EVs and ^CTX^RBC-EVs (hereafter referred to as RBC-EV_cispt_ and ^CTX^RBC-EV_cispt_, respectively) were rinsed to 1mL with PBS 1X without CaCl_2_ and MgCl_2_ (Corning, NY, USA) and ultracentrifuged at 100000 x*g* for 2h (Optima MAX-XP centrifuge equipped with a TLA-55 rotor, Beckman Coulter, USA). The amount of platinum in RBC-EV_cispt_ was measured by inductively coupled plasma mass spectrometry (ICP-MS) Elan DRC II (Perkin Elmer, Waltham, MA, USA) employing the external calibration quantitative analysis method. The instrument was calibrated with a calibration curve of different platinum concentrations of 1-5-10 ppb, starting from the smart solutions O_2_Si Platinum standard solution at 1000 ppm in 5% HCl. Samples were diluted 1:1000 in distilled water. The intra-series coefficient of variation ranged between 4% and 8%, while the inter-series coefficient of variation varied between 2% and 6%. The detection limit was calculated with a 3x standard deviation of the blank and was established as 0.05 ppb. The measurement range spanned from 0.1 ppb to 500 ppb. Cisplatin concentration was calculated as Platinum concentration (g/L) x cisplatin molecular weight (g/mol) / Platinum molecular weight (g/mol). For the optimization of the protocol for cisplatin encapsulation, results from six different preparations were used. For the comparison of cisplatin encapsulation between ^CTX^RBC-EV_cispt_ and RBC-EV_cispt_, results from four different preparations were used and expressed as mean ± SEM.

### Cisplatin release kinetics assay

RBC-EV_cispt_ were obtained by incubating RBC-EVs and cisplatin (1.4 mg/mL) for 24 h at 37°C as mentioned above. RBC-EV_cispt_ pellets were resuspended in 200 µL of PBS 1X without CaCl_2_ and MgCl_2_. 40 µL were transferred in another tube and stored for platinum quantification, while the remaining 160 µL were loaded into a 12-14kDa Pur-A-Lyzer dialysis column (Sigma-Aldrich, St. Louis, MO, USA) and dialyzed against 1mL of PBS at 37°C. 100µL of the outer buffer were collected at 0, 2h, 4h, 10h and 24h. At each time point, 100 µL of fresh PBS 1X was added. The platinum amount was quantified by ICP-MS and cisplatin concentration was calculated as mentioned above. The percentage of cisplatin released was obtained, calculating at first the total mg in 1mL for each time point. This amount was then summed to the total mg of the previous time point. The percentage point by point was then normalized on the initial concentration of cisplatin.

### Cell line and culture conditions

Human Triple-Negative Breast Cancer (TNBC) MDA-MB-231 cells (HTB-26, ATCC, Manassas, VA, USA) and the Her2-enriched Human Breast Cancer BT-474 cell lines were cultured in Dulbecco’s Modified Eagle Medium with 4.5 g/L glucose, L-glutamine and sodium pyruvate (DMEM, Corning, USA), supplemented with 10% (v/v) fetal bovine serum (FBS, Immunological Science, UK), 1% (v/v) penicillin/streptomycin (Corning, USA), and hereafter referred to as complete media. Cells were maintained in a humidified 5% CO_2_ incubator at 37°C and routinely tested for mycoplasma contamination using the MycoBlue Mycoplasma Detection kit (Vazyme, Nanjing, China).

### Patient-derived organoids establishment

Two TNBC patients (patient #1 and patient #2) enrolled at the Breast Unit of ICS Maugeri IRCCS (Pavia, Italy) in the protocol of “Bruno Boerci Oncological Biobank” (ICS Maugeri IRCCS’s ethical committee approbation of 27 July 2009) between April 2019 and July 2020, were a 45- and a 60-years-old women, respectively. Patient #1 displayed an invasive ductal carcinoma of over 6 cm in right mammary gland and reported familiarity for breast and ovarian neoplasia, as confirmed by the presence of BRCA1 gene mutation. Patient #1 has been subjected to neoadjuvant chemotherapy (NAC) with 4 cycles of Epirubicin and Cyclophosphamide, and 12 cycles of paclitaxel and carboplatin followed by radical mastectomy. During surgery, a biopsy specimen was collected, analysed and used to establish PDO #1 culture. Pathological evaluations performed on this specimen confirmed the classification as invasive ductal carcinoma, with a TNBC molecular subtype (Estrogen 0%, Progesterone 0%, Ki67 60%, c-erbB2 1+). Patient #2 displayed a lesion of 21 mm classified as invasive ductal carcinoma, TNBC molecular subtype (Estrogen 0%, Progesterone 0%, Ki67 70%, c-erbB2 0) in right mammary gland and reported familiarity for breast and ovarian neoplasia, despite only the presence of a BRCA1 gene variant with unknown significance has been detected. During clip positioning, another biopsy specimen was collected, analysed and used to establish PDO #2 culture. The bioptic samples from patient #1 and #2 have been collected in Ad-DF +++ medium (Hyclone DMEM-F/12 1:1 supplemented with 10 mM HEPES, 1% Penicillin/Streptomycin and 1% L-glutamine) and stored at 4°C until processing. Then, samples were cut into 1-3 mm^3^ and two random pieces were fixed in formalin and embedded in paraffin to perform Hematoxylin and Eosin staining (H&E) and Immunohistochemistry (IHC) labelling by using routine procedures. Primary PDO #1 and PDO #2 cultures were obtained following the procedure described in our consolidated protocol (31). Briefly, the collected tissue was removed of the adipose tissue and mechanically and enzymatically digested in 10 mL Ad-DF +++ medium supplemented with 500 µL of Collagenase (20 mg/mL) and 10 µL of Y27632 ROCK inhibitor (10 mM) for 1-2 h at 37°C. The sample was filtered to remove the undigested tissue, collected in a 15 mL tube and centrifuged. The pellet was washed twice and then resuspended in the appropriate amount of basement membrane extracts (BME) and seeded in a pre-warmed multi-well plate. Once BME-PDO drops were solidified, the appropriate amount of culture medium (CM; DMEM/F12 1×, L-glutamine 1%, Penicillin/Streptomycin 1%, Hepes 10mM, Noggin conditioned medium 25×, B27 supplement 1×, N-acetyl-cysteine 1.25 mM, Nicotinamide 0.2 mM, A 83-01 500 nM, Y-27632 5 µM, R-spondin1 conditioned medium 10%, Primocin 50 µg/mL, Human EGF 5 ng/mL, FGF-10 Human recombinant 20 ng/mL, KGF/FGF-7 Human recombinant 5 ng/mL, Heregulin-beta-1 Human recombinant 37.5 ng/mL, SB 202190 500 nM) was added depending on the multi-well size and changed every 2-3 days. Every 7-10 days, when confluence was achieved, PDO #1 and PDO #2 cultures were collected and passed. Each organoid culture was frozen in Cell Culture Freezing medium (Gibco, Thermo Fisher Scientific, Waltham, MA, USA) and transferred to liquid nitrogen for long-term storage.

### Patient-derived organoids characterization

BME-organoid drops were removed with a sterile cell lifter from the plate and transferred into a mould containing a layer of optimal cutting temperature compound (OCT). Once the drops were included in OCT, the mould was kept at -80°C until processing. The OCT-embedded PDOs were sectioned to obtain histological slices of about 3 µm thicknesses. After the fixation, the histological slides were stained with H&E, labelled with VENTANA BenchMark ULTRA (Roche Diagnostics, Monza, Italy), following automatized IHC protocols for Estrogen receptor, Progesterone receptor, c-ErbB2 (i.e. HER2) and Ki67 (32). For the comparison between organoids and the tumour of origin, two random pieces of the surgical tissues were fixed in formalin and embedded in paraffin to perform H&E and IHC labelling by using routine procedures. For morphological Transmission Electron Microscopy (TEM) and Scanning Electron Microscopy (SEM) analysis, PDO #1 and PDO #2 were processed and analysed as previously described (31). For immunofluorescence analysis, 3×10^6^ organoids were isolated from BME after treatment with Dispase (1 µg/mL at 37°C for 1-2 h). Then, PDOs were collected, washed in PBS thrice and fixed with Paraformaldehyde (PFA) 4% for 15 minutes at room temperature (RT). After the fixation, PDOs were washed in PBS thrice and then permeabilized using Triton X-100 0.1% for 10 minutes at RT. After three washes, PDO pellets were resuspended in 500 µL of blocking solution containing 2% goat serum and 2% Bovine Serum Albumin (BSA) in PBS 1X for 1 hour at RT. PDOs were incubated with the primary antibodies in blocking solution for 2 h at RT. We used the following primary rabbit antibodies directed to Ki67 (Abcam ab243878, 1:500), EGFR (Genetex GTX35199, 1:200), Vimentin (Genetex GTX100619, 1:500) and HER-2 (Cell Signalling Technologies #2165, 1:200) proteins. PDOs were washed thrice in PBS and incubated with the secondary antibodies anti-Rabbit Alexa Fluor 546 (1:300), Wheat Germ Agglutinin FITC (1:300), DAPI (1:10000), in blocking solution overnight at 4°C. After the staining, PDOs were washed three times in PBS 1× and seeded on specimen slide in mounting medium ProLong^TM^ Gold (Invitrogen, P36935, Thermo Fisher Scientific) for the acquisition with the Leica confocal microscope SP8 equipped with 405, 488 and 513 nm lasers. Acquisition was performed at 1024×1024 dpi resolution. For CD24, CD44, CD49f and EPCAM staining, 3×10^6^ organoids were isolated from BME by incubating them with Dispase 1 µg/mL. Once collected, PDOs were reduced into single cells through the shearing procedure using TrypLe^TM^ Select (1x; Gibco, 12563-029). After three washes with Hank’s Balanced Salt Solution (HBSS from HyClone, SH30268.02), the cells were fixed with PFA 4% for 5-10 minutes in ice. Fixed cells were washed thrice with HBSS supplemented with FBS 2% and aliquoted into four tubes containing about 7.5 × 10^5^ cells each. The first tube was labelled with the Lineage PE cocktail of antibodies (PE mouse anti-human CD2, Cod. 555327, 1:100; PE mouse anti-human CD3, Cod. 555333, 1:00; PE mouse anti-human CD10, Cod. 555375, 1:100; PE mouse anti-human CD16, Cod.555407, 1:100; PE mouse anti-human CD18, Cod. 555924, 1:100; PE mouse anti-human CD31, Cod. 555446, 1:100; PE mouse anti-human CD64, Cod. 558592, 1:100; PE mouse anti-human CD140b, Cod. 558821, 1:100; BD Biosciences) for 15 min at RT, to set the gate of lineage positive cells, which should be excluded from the analysis. The second tube contains only unstained cells, to acquire negative signals. A third tube was labelled 15 min at RT with lineage cocktail FITC mouse anti-human CD24 (Cod. 555427, 1:50, BD Biosciences) and APC mouse anti-human CD44 (Cod. 559942, 1:50, BD Biosciences), to identify CD24/CD44 cell population, while the fourth tube was labelled with lineage cocktail supplemented with FITC rat anti-human CD49f (Cod. 555735, 1:50, BD Biosciences) and APC mouse anti-human EPCAM (Cod. 347200, 1:100, BD Biosciences), to identify CD49f/EPCAM populations. After the staining, labelled cells were washed thrice with HBSS supplemented with FBS 2% and analysed using CytoFLEX flow cytometer (Beckman Coulter). Acquisition was performed on 20000 events, within the selected region of singlets viable cells. For EGFR evaluation, 1.5×10^6^ organoids were isolated from BME, reduced into single cells and fixed with 4% paraformaldehyde (PFA, 76240, Thermo Fisher Scientific, USA) for 5 minutes at 4°C. Cells were washed thrice with HBSS supplemented with FBS 2% and were transferred into two tubes containing about 7.5×10^5^ cells for each. A tube was labelled with the primary chimeric monoclonal antibody Cetuximab (CTX, 1:200) for 15 min at RT. Labelled cells were washed thrice with HBSS supplemented with FBS 2%. Both tubes were labelled with the AlexaFluor488 (AF488) goat anti-Human secondary antibody (Thermo Fisher, 1:300). The tube containing cells labelled with only the secondary antibody was used to set the region of positivity. After the staining, labelled cells were washed thrice with HBSS supplemented with FBS 2% and analysed as described above.

### Labelling of RBC-EV preparations

RBC-EV preparations were labelled with MemGlow™ 488 (MG01-10, Cytoskeleton Inc., Denver, CO, USA) as previously described (27). Briefly, RBC-EV samples were incubated with 100 nM MemGlow™ 488 for 30 minutes at RT. Excess of free probe was removed by ultracentrifugation at 100000 xg for 2 h at 4°C (Optima MAX-XP equipped with a TLA-55 rotor, Beckman Coulter, USA).

### Assessment of EGFR expression on MDA-MB-231 cells

2×10^6^ MDA-MB-231 cells and 2×10^6^ of BT-474 cells (used as control for low EGFR expression) were harvested and fixed with 4% paraformaldehyde (PFA, 76240, Thermo Fisher Scientific, USA) for 5 minutes at 4°C. Cells were washed in PBS for three times and the pellet was resuspended in PBS, 2% Bovine Serum Albumin (BSA, A2153, Sigma-Aldrich, USA), 2% goat serum (G9023, Sigma-Alrdich, USA). About 5×10^5^ of fixed cells were transferred in tubes and immunodecorated with the anti-EGFR antibody (Cetuximab, CTX, 1 µg/tube;) PBS, 2% BSA; Sigma) and 2% goat serum (Euroclone) for 30 min at RT. Then, cells were washed thrice with PBS and immunodecorated with Alexa Fluor 488 goat anti-human secondary antibody (1 µL/tube; Thermo Fischer Scientific) in PBS, 2% BSA and 2% goat serum for 30 min at RT. After three washes with PBS, cells (n=3) were analyzed by CytoFLEX flow cytometer (Beckman Coulter). 20000 events were acquired for each analysis, after gating on viable cells and on singlets. A sample of cells immunodecorated with the secondary antibody only was used to set the region of positivity.

### Cellular RBC-EV uptake studies in 2D cell culture system by confocal imaging

For imaging analysis, MDA-MB-231 cells were seeded onto 12 mm-sized coverslips pre-coated with 50 µg/mL collagen (CliniSciences, Nanterre, France) as 30000 cells/coverslip (26549 cells/cm^2^) placed in 12-well tissue culture plates (Euroclone, Italy). After 24h, cells were treated MemGlow488-labelled-RBC-EVs, ^CTX^RBC-EVs, RBC-EV_cispt_, and ^CTX^RBC-EV_cispt_ for 4h and 24h in 1 mL of complete media. For the investigation of the comparison of RBC-EVs and ^CTX^RBC-EVs uptake, the following RBC-EV and CTX final concentrations were used: 6.2x10^10^ RBC-EVs (corresponding to 1.8x10^10^ EVs/cm^2^) and 35 nM CTX. For the comparison of RBC-EV_cispt_ and ^CTX^RBC-EV_cispt_ uptake, the followings: 7.3x10^10^ RBC-EVs (corresponding to 2.7x10^10^ EVs/cm^2^), 5µM cisplatin and 22 nM CTX. Cells were washed twice with PBS without CaCl_2_ and MgCl_2_ (Corning, USA) and fixed with a 3% paraformaldehyde (PFA, Thermo Fisher Scientific, USA) solution for 15 min at RT. PFA was quenched with 50 mM NH_4_Cl for 10 minutes at RT. The cells were washed twice with PBS 1X and permeabilized with 0.3% saponin (Thermo Fisher Scientific, USA) in PBS 1X (PBS-S) for 10 minutes at RT. Intracellular hemoglobin was stained with anti-HBB antibody diluted 1:50 (H00003043-M02A, Abnova, Taiwan) in PBS-S for 1 hour at RT. The coverslips were washed twice with PBS-S and incubated with the AlexaFluor 647-secondary antibody Rabbit anti-Mouse diluted 1:400 (IS20285-1, Immunological Sciences, Rome, Italy) and DAPI diluted 1:600 in PBS-S (D3571, Invitrogen, USA) for 30 minutes at 37°C. Coverslips were washed twice with PBS 1X and once with double-distilled water (ddH_2_O), and finally mounted with ProLong™ Gold Antifade Mountant (P36934, Invitrogen, USA). 1024 × 1024 pixel images were acquired with a Zeiss LSM 900 confocal microscope with a Plan-Apochromat 63×/1.4 Oil DIC or EC Plan-Neofluar 40x/1.3 Oil DIC objectives, with a 0.2% 405 nm laser for DAPI, 0.2% 488 nm laser for MemGlow™ 488 and 0.4% 633 nm laser for AlexaFluor 647-secondary antibody.

### Cell binding assay

5×10^5^ MDA-MB-231 cells were incubated for 2 h at 4°C in PBS, 2% BSA and 2% goat serum (Euroclone) supplemented with 10^9^ or 10^10^ RBC-EVs/mL, 10^9^ or 10^10^ ^CTX^RBC-EVs/mL or CTX free (2.43nM or 24.3 nM) in 5 mL tubes. RBC-EVs were fluorescently labelled with Memglow88, while CTX free or conjugated to RBC-EVs were labelled with Cy7.5, as previously described. Then, cells were washed thrice with PBS and cells were analyzed by CytoFLEX flow cytometer (Beckman Coulter). 20000 events were acquired for each analysis, after gating on viable cells and on singlets. Samples of untreated cells were used to set the regions of positivity.

### Competition assay

5×10^5^ MDA-MB-231 cells/sample were incubated for 1 h at 4°C with 2.8 µM of free unlabelled CTX as competitor (1 mL) in PBS supplemented with 0.3% BSA. Then, cells were centrifuged to remove unbound CTX and incubated with Cy7.5-labelled CTX or Cy7.5-labelled ^CTX^RBC-EVs for 1h at 4°C. At the end of incubation, cells were washed thrice, resuspended in PBS (500 µL) and analyzed by CytoFLEX flow cytometer (Beckman Coulter). 20000 events were acquired for each analysis, after gating on viable cells and on singlets. A sample of untreated cells was used to set the appropriate gates.

### Cellular cisplatin uptake studies in 2D and 3D cell culture systems

For cisplatin uptake analysis in 2D culture conditions, MDA-MB-231 cells were seeded into collagen-precoated 35 mm tissue culture dishes (ET2035, Euroclone, Italy) as 22500 cells/cm^2^. After 24h, cells were washed once with PBS 1X without CaCl_2_ and MgCl_2_ and treated for 30 min and 24h with RBC-EV_cispt,_ ^CTX^RBC-EV_cispt_, and free cisplatin in 2 mL of complete media. Treatments were carried out using the following final concentrations: 5.6 x 10^10^ RBC-EVs/mL (corresponding to 1.4x10^10^ EVs/cm^2^), 5µM cisplatin and 11.5 nM CTX. Cells were washed once with PBS 1X without CaCl_2_ and MgCl_2_ and harvested using 0.25% trypsin-EDTA solution (Corning, USA). Cells were counted with a Burker chamber and centrifuged twice at 800 xg for 10 minutes at 4°C (Eppendorf, Germany). The obtained pellets were digested with 200 µL of HNO_3_ and 200 µL of ddH_2_O and incubated at 70°C for 1h. The digested pellets were diluted 1:50 and measured with ICP-MS as previously described. Data from three independent experiments are expressed as mean ± SEM.

### Subcellular fractionation

MDA-MB-231 cells were seeded into collagen-precoated 35 mm tissue culture dishes as 22500 cells/cm^2^. After 24h, cells were washed once with PBS 1X without CaCl_2_ and MgCl_2_ and treated in triplicate for 24h with RBC-EV_cispt_, cisplatin and ^CTX^RBC-EV_cispt_. Treatments were carried out using the following final concentrations (final volume of 2 mL): 7.3 x 10^10^ EVs/mL (corresponding to 1.8x10^10^ EVs/cm^2^), 5µM cisplatin and 19.5 nM CTX. Cellular fractioning was carried out following the procedure previously described (33). Briefly, cells were washed twice with cold PBS 1X without CaCl_2_ and MgCl_2_, scraped and spinned for 10 seconds at 4°C. Pellets were resuspended in 100 µL of PBS 1X containing 0.1% NP-40 (PanReac Applichem, USA). 20µL of the whole homogenate was used for Western Blot analysis. Samples were further centrifuged for 10 seconds and 80 µL of the supernatant, corresponding to the cytoplasm fraction, was transferred into a new tube: 50µL were used for ICP-MS measurements, while 30 µL for Western Blot analysis. Pellets were washed once with PBS 1X containing 0.1% NP-40 and spinned down for 10 seconds. Pellets, corresponding to the nuclear fraction, were resuspended in 80 µL of PBS 1X containing 0.1% NP-40: 50µL were used for ICP-MS measurements, while 30 µL for Western Blot analysis. For ICP-MS, the platinum content was measured by diluting the sample 1:50 and digested with 200 µL of HNO_3_ and 200 µL of ddH_2_O. Samples were incubated at 70°C for 1h and diluted 1:50 with ddH_2_O. Data are expressed as mean ± SD. For Western Blot analysis, fractions were heated in 6X Laemmli buffer and separated by SDS–PAGE on acrylamide/bisacrylamide gels and then transferred onto a PVDF membrane as described above. The following secondary antibodies were used: α-LAMP-1 (1:500, Clone H4A3, sc200-11, Santa Cruz Biotechnology Inc., Dallas, TX, USA) for the cytoplasmatic fraction, α-lamin A/C (1:1000, mab636, Thermo Fisher Scientific, USA) for the nuclear fraction and anti-HBB (1:500, H00003043-M02A, Abnova, Taiwan) to confirm hemoglobin uptake.

### 3-(4,5-dimethylthiazol-2-yl)-2,5-diphenyl-2H-tetrazolium bromide (MTT) assay

MDA-MB-231 cells were seeded in 96-well plates in complete media as 15000 cells/cm^2^. After 24h, cells were treated in quadruplicate for 4h and 24h with RBC-EV preparations, cisplatin and CTX. Treatments were carried out using the following average concentrations in 0.1 mL: 6.2 x 10^10^ EVs/mL (corresponding to 1.94 x 10^10^ EVs/cm^2^), 5µM cisplatin and 15 nM CTX. After the treatments, 0.01mL of the 5 mg/mL MTT solution (M2128-1G, Sigma-Aldrich, USA) was added and the cells were further incubated for 4h. The formazan crystals were dissolved with 0.1 mL of DMSO (276855, Sigma-Aldrich, USA) and the absorbance was measured with Ensight MultiMode Reader (Perkin Elmer) at 595 nm. The absorbance values of the blank solution (DMSO) were removed from each sample. Cell viability percentage was calculated as follows: (Abs_595_ treated cells/ Abs_595_ untreated cells) x 100. Data from three independent experiments are expressed as mean ± SEM. The dose- and time-dependent cytotoxicity assessment of all the building blocks (namely, cisplatin and cetuximab as free drugs and RBC-EVs) were performed at the reported concentrations and time as mentioned above.

### Measurement of oxygen consumption and extracellular acidification rates

The rate of change of dissolved O_2_ (oxygen consumption rate, OCR) and glycolysis rate (as extracellular acidification rate, ECAR) were measured with the Seahorse XFe24 Extracellular Flux Analyzer (Agilent, Santa Clara, CA, USA). MDA-MB-231 cells were seeded in 100 μL in XFe24 V7 PS Cell Culture Microplates (Agilent). The plates were incubated for 1 hour at RT, and then for 5 h in 5% CO_2_ atmosphere at 37°C to optimize cell adhesion. The medium volume was brought up to 250 μL and then microplates were incubated in 5% CO_2_ atmosphere at 37°C for 24h. Thereafter, cells were treated in quadruplicate for 24h with RBC-EVs, CTX, and cisplatin alone or in combination, diluted in fresh complete growing medium. After treatment, the medium was replaced with 500 uL of Seahorse XF DMEM medium pH 7.4 (Agilent) supplemented with 10 mM glucose, 2 mM L-glutamine, and 1mM sodium pyruvate. Microplates were incubated for 1h at 37°C w/o CO_2_. At the end of the incubation, both OCR and ECAR were measured with the Seahorse XF Cell Mito Stress test (Agilent). After basal OCR measurement, Oligomycin (2 µM), Carbonyl cyanide-4-(trifluoromethoxy) phenylhydrazone (FCCP, 0.5 µM) and a mixture of Rotenone and Antimycin A (0.5 µM each) were sequentially injected to monitor uncoupled (proton leak), maximal and non-mitochondrial respiration, respectively. Data were normalized to DNA content using the CyQUANT^TM^ Cell Proliferation Assay Kit (Invitrogen), according to the manufacturer’s guidelines. Data analysis and calculation of ATP-linked respirations and spare respiratory capacity were performed with the Seahorse Wave software (version 2.6.3, Agilent).

### RNA-seq library preparation and bioinformatic data analysis

MDA-MB-231 cells were seeded into 35 mm tissue culture dishes as 22500 cells/cm^2^. After 24h, cells were treated for 24h with 4.6 x 10^10^ RBC-EVs/mL, 4.6 x 10^10^ RBC-EV_cispt_/mL (5µM cisplatin), 4.6 x 10^10^ ^CTX^RBC-EV_cispt_/mL (5µM cisplatin e 7.7 nM CTX), 4.6 x 10^10^ ^CTX^RBC-EVs/mL (7.7 nM Cetuximab), free CTX (7.7 nM), free cisplatin (5 µM), and a mix of CTX (7.7 nM) + Cisplatin (5 µM), in 2mL for each treatment. Each treatment was carried out in triplicate. Then, cells were detached using 0.25% trypsin-EDTA solution, washed in PBS 1X and centrifuged at 1000 xg for 5min. Pellets were snap frozen in dry ice and stored at -80°C until use. For each treatment condition, total RNA was extracted by using RNeasy Mini Kit (Qiagen, Hilden, Germany), according to manufacturer’s protocol, and treated with RNase-free DNAse I (Thermo Fisher Scientific, USA). RNA samples were checked for quality on Agilent 2200 TapeStation System (Agilent Technologies, Santa Clara, CA, USA) and quantified by Qubit 4 Fluorometer using the Qubit RNA HS Assay Kit (Thermo Fisher, Invitrogen). Starting from 400 ng of each isolated RNA, RNA-seq libraries were prepared using the Illumina TruSeq Stranded mRNA Library Prep Kit (Illumina, San Diego, CA, USA), according to manufacturer’s instructions. Each library was checked on Agilent 2200 TapeStation System (Agilent Technologies) and quantified by Qubit 4 Fluorometer using the Qubit DNA HS Assay Kit (Thermo Fisher). Sequencing was carried out in 2×150-cycle runs on Illumina HiSeq4000 platform. Three independent biological replicates were prepared and sequenced for each condition. After fastq quality control by using FastQC tool (v.0.11.8), raw reads were trimmed to 100 bases and mapped to the human reference genome using STAR aligner (v.2.7.10a) (34). Gene counts were calculated by HTSeq package (v.0.11.1) (35) using the hg38 Encode-Gencode GTF file (v39) as gene annotation file. Differential gene expression analysis was carried out using DESeq2 Bioconductor/R package (v.1.30.1). RUVSeq R package (v.1.24.0) (36) was used for batch effect removal (37). Low expressed genes (sum of read counts across all samples < 10), as well as outlier samples, were filtered out before testing genes for statistical significance. A |log2FC| > 1 and adjusted p-value (padj, Benjamini-Hochberg (BH) correction) < 0.1 were used as cut-off to define statistically significant differentially expressed genes (DEGs). ClusterProfiler R package (v.4.6.2) was used to perform functional enrichment over-representation analysis (ORA) on Gene Ontology Biological Process (GO-BP) categories and biological pathway collections (KEGG, Reactome, WikiPathways, MSigDB hallmark gene sets (38)) false discovery rate (FDR, BH correction) < 0.05 was applied to all the annotation terms to define statistically significant enrichments. Raw sequence data are available in NCBI Short Reads Archive (SRA) under Accession Number PRJNA1067532.

### 3D cytotoxicity assay

To establish a cells viability assay, the organoids were sheared 2-3 days before the seeding to obtain smaller and uniform in size PDOs. The organoids were isolated from BME by Dispase treatment, collected in 15 mL tube and washed twice with Ad-DF +++. The PDOs were counted, diluted in CM containing 10% BME and seeded 10000 cells/well in a 96-wells spheroid microplate (Corning) at the concentration of 200 cells/µL. After 24 h, four different concentrations of Cetuximab (Erbitux® 5mg/mL, Merck) and of Cisplatin (1 mg/mL, Sandoz) (both ranging from 0.5 nM to 200 nM) were added in 10 replicates. Untreated cells were used as negative control. After three days of expansion at 37°C and 5% CO_2_, the Cell Titer Glo 3D Kit (Promega) was used, to measure the ATP content as an indicator of cell viability, according to manufacturer’s instructions. Emitted luminescence was read in microplate reader (PerkinElmer, Victor Nivo Multimode) and data were analysed using GraphPad Prism 8 software (GraphPad Software Inc., La Jolla, CA, USA).

### Ex vivo cisplatin toxicity assessment on red blood cell morphology

RBC concentrates were extracted from a healthy donor (0+) stored in Saline Adenine Glucose Mannitol (SAGM) at 4°C. After 24h, 6.23x10^9^ RBCs were incubated with cisplatin (5µM and 25µM), 5.5x10^10^ ^CTX^RBC-EV_cispt_/mL (5µM cisplatin, 20 nM CTX) for 3 h in 1mL final volume. After incubation, the blood smear was performed, and RBCs were fixed on the glass slides in 100% ethanol for 5 minutes. PBS was used as vehicle. RBC morphologies after treatments were examined at 20x magnification with a Axio Vert microscope (Zeiss, Jena, Germany). Cell counting was performed with Fiji software and the relative percentage of RBC morphologic alterations were calculated as follow: (Number of RBCs with a specific morphology/Total counted RBCs) x 100. Data are expressed as mean ± SD.

### Lipid peroxidation detection with C11-BODIPY^581/591^ on MDA-MB-231 cells

For lipid peroxidation assessment in 2D culture conditions MDA-MB-231 cells were seeded in triplicate onto 12 mm-sized coverslips pre-coated with 50 µg/mL collagen (CliniSciences, Nanterre, France) as 50 000 cells/coverslip (44247 cells/cm^2^) placed in 12-well tissue culture plates (Euroclone, Italy). After 24h, cells were treated for 24h with 4.6 x 10^10^ RBC-EVs/mL, 4.6 x 10^10^ RBC-EV_cispt_/mL (5µM cisplatin), 4.6 x 10^10^ ^CTX^RBC-EV_cispt_/mL (5µM cisplatin e 7.7 nM CTX), 4.6 x 10^10^ ^CTX^RBC-EVs/mL (7.7 nM Cetuximab), free CTX (7.7 nM), free cisplatin (5 µM), and a mix of CTX (7.7 nM) + Cisplatin (5 µM) in 2mL for each treatment. As negative control, cells were added with fresh cell culture medium (untreated), as positive control, cells were treated with Erastin 5 µM for 1 h. All treatments were conducted in complete cell culture medium as described above. After treatments cell culture medium was replaced with fresh one and cells were incubated with BODIPY™ 581/591 C11 5 µM for 40 minutes (MedChemExpress). Cells were then fixed and stained as previously described (39). Briefly, cells were washed with PBS 1X and fixed with 3% paraformaldehyde (PFA) for 15 min, washed with NH_4_Cl for 15 min. Nuclei were stained with DAPI (diluted 1:600 in PBS) for 30 minutes. Coverslips were washed twice with PBS 1X and once with double-distilled water (ddH_2_O), and finally mounted with ProLong™ Gold Antifade Mountant (P36934, Invitrogen, USA). Detection of lipid peroxidation was performed by fluorescent microscopy(40,41): briefly, 1024 × 1024 pixel images were acquired with a Zeiss LSM 900 confocal microscope with a EC Plan-Neofluar 40x/1.3 Oil DIC objectives, with a 1.2% 405 nm laser for DAPI, 2.2% 488 nm laser and 1.6% 561 nm laser for C11-BODIPY^581/591^. The green and red fluorescence of C11-BODIPY^581/591^ was acquired simultaneously using double wavelength excitation (laserlines 488 and 561 nm) and detection (emission bandpass filters 530/30 and 590/30) (41,42). Fluorescence intensities in the red and green channels were quantified using Fiji software. For each channel, a fluorescence threshold was established based on untreated cells. Green and red fluorescence intensities from all treatment conditions were normalized to the corresponding untreated thresholds. The green-to-red fluorescence intensity ratio was calculated as an oxidation index to quantify lipid peroxidation (43). Twenty-one images were analyzed for each treatment and for the positive control.

### Statistical analysis

For cisplatin loading optimization, statistical analysis was performed using GraphPad Prism 8.0a version software with an unpaired t-test, with a two-sided p-value calculations (95% confidence interval, statistical significance when *p* < 0.05). For MDA-MB-231 cytotoxicity screening and cisplatin uptake, statistical analysis was performed using the ordinary one-way ANOVA with Tukey’s multiple comparison tests, with a single pooled variance. For the dose- and time-dependent cytotoxicity assessment of all the building blocks, statistical analysis was performed using the ordinary two-way ANOVA with Tukey’s multiple comparison test. For EGFR expression, statistical significance was determined by two-tailed unpaired Student’s t-test with Welch’s correction. For binding and competition assays, significance was determined by ordinary one-way ANOVA corrected with Sidak’s multiple comparison tests. Drug screening results have been analyzed by ordinary one-way ANOVA performing the Dunnett’s multiple comparison test, analyzing both the effect of drug and organoid type. Statistics was evaluated using GraphPad Prism 8.0a version (GraphPad Software Inc., La Jolla, CA, USA). Data are reported as mean ± Standard Error Mean (SEM). The level of statistical significance was set at p = 0.05.

For lipid peroxidation analysis ordinary one-way ANOVA performing the Dunnett’s multiple comparison test was conducted. Statistics was evaluated using GraphPad Prism 8.0a version (GraphPad Software Inc., La Jolla, CA, USA). The level of statistical significance was set at p = 0.05.

## Results and Discussion

### Cisplatin encapsulates in RBC-EVs *via* co-incubation

Red blood cell-derived EVs (RBC-EVs) were produced upon induction with calcium and calcium ionophore and characterized according to their biochemical and biophysical properties, following MISEV2018 and MISEV2023 guidelines (44,45). RBC-EVs were enriched in typical RBC protein markers, such as Band 3 anion transport protein, hemoglobin (subunit β, HBb), and the EV marker flotillin-1 (Figure S1A). As shown earlier by our studies, RBC-EVs produced with our induction method contain other typical EV markers as Annexin XI and Adam10, but do not contain classical EV markers as tetraspanins, aligning with literature data (24,46,47). The Colorimetric NANoplasmonic (CONAN) assay (30) revealed that RBC-EV preparations contained a negligible amount of soluble non-EV associated proteins (SAPs), and therefore can be considered pure from free contaminants (aggregation index (AI) ≤ 20%, Figure S1B). From ≈300 mL of RBC solution, we obtained RBC-EV preparations with an average protein concentration of 703 ± 123 µg/mL, while Nanoparticle Tracking Analysis (NTA) confirmed the presence of 2.4 x 10^12^ ± 5.0 x 10^10^ EVs/mL, with a mean diameter of 193 ± 7.3 nm (Figure S1C, D). Subsequently, we optimized cisplatin loading into RBC-EVs by a co-incubation reaction at 37°C for both 4 h and 24 h in agitation, by adapting protocols previously described for other hydrophilic drugs (48–51) (Figure 1A). Cisplatin loading yield *per* EVs was calculated by Inductively Coupled-Plasma Mass Spectrometry (ICP-MS). We obtained a statistically significant 3-fold increase in cisplatin loaded amount after 24h co-incubation as compared to the 4h co-incubation (4.4 x 10^-14^ µmol cis/EV for 4h *vs.* 1.4 x 10^-13^ µmol cis/EVs for 24h co-incubation) with a loading efficiency of 0.91% +/- 0.2. (Figure 1B). Considering this, all the subsequent experiments were performed with RBC-EV_cispt_ after 24 h incubation. We compared RBC-EVs and RBC-EV_cispt_ morphology and size distributions by Atomic Force Microscopy (AFM) imaging and NTA, respectively. AFM imaging revealed the presence of round-shaped nanoparticles in both RBC-EV samples and the absence of broken EVs or lipid-debris in RBC-EV_cispt_ preparation (Figure 1C). Both RBC-EVs and RBC-EV_cispt_ showed similar size distributions (Figure 1D), although we observed a fraction of bigger nanoparticles (NPs) in RBC-EV_cispt_ formulation and the mean EV diameter shifted from 193 ± 7.3 nm for RBC-EVs to 218 ± 9.0 nm for RBC-EV_cispt_ (Figure S1D). This size shift for RBC-EV_cispt_ could be due to the process of cisplatin loading which might cause restructuring of the lipid bilayer (i.e. rearrangement of lipids), leading to vesicle shape changes and, consequently, larger apparent size (48–51).

**Figure 1.**
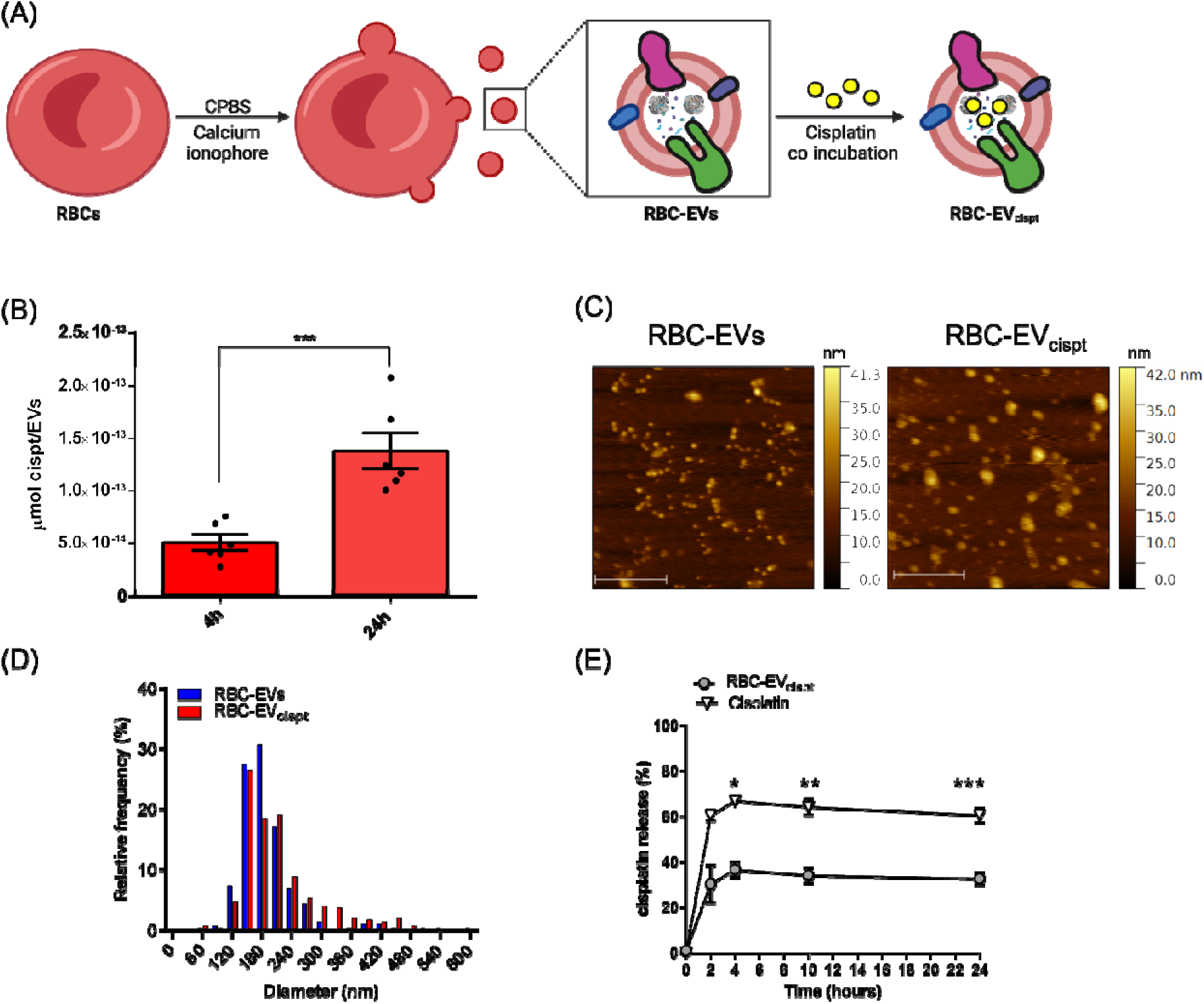
Characterization of cisplatin-loaded RBC-EVs. **(A)** Scheme on the production of RBC-EV_cispt_ *via* co-incubation method carried out at 37°C for 4 and 24 h. **(B)** Comparison between the 4h- and 24h-incubation in RBC-EV_cispt_ in terms of cisplatin encapsulation yield. Unpaired t-test: *p* < 0.05, \*\*\**p* = 0.0008. **(C)** Representative AFM images of RBC-EVs and RBC-EV_cispt_ after 24 h co-incubation with cisplatin. Scale bar: 1µm. **(D)** Size distributions of RBC-EVs and RBC-EV_cispt_ after 24 h co-incubation with cisplatin obtained with NTA measurements. **(E)** Cisplatin release pattern from cisplatin solution and RBC-EV_cispt_. Results are expressed as mean (n=3) ± SD. Paired t-test: *p = 0.0185, ** = 0.0017, *** 0 0.0008.

Given cisplatin ability to passively cross biological membranes, we measured the drug release from free cisplatin solution and RBC-EV_cispt_ using a 10kDa cut-off dialysis membrane. Cisplatin was released in a burst-release manner after the first 2 h for both free cisplatin (up to 60%) and RBC-EV_cispt_ (up to 20%), and then the trend slowed. (Figure 1E). After 2h, cisplatin was released with a constant rate, suggesting a controlled release profile from RBC-EV_cispt_ (Figure 1E). Since it has already been reported that cisplatin as well as other platinum complexes can strongly interact with hemoglobin, we cannot exclude that part of the encapsulated cisplatin could be also retained inside the RBC-EV_cispt_, probably thanks to its binding with luminal proteins (e.g., Hb) (52). Further investigations will be needed to characterize the mode of cisplatin association with RBC-EVs, specifically to determine whether the drug is encapsulated within the EVs or adsorbed onto their surface.

### Engineering RBC-EVs with cetuximab confers a higher uptake in MDA-MB-231 cell line

In parallel with cisplatin encapsulation, we set up RBC-EV functionalization with the CTX monoclonal antibody, as sketched in Figure 2A. CTX was added onto RBC-EV membrane (^CTX^RBC-EVs) using a bio-orthogonal click chemistry approach, as we have previously described (27). The functionalization process led to a yield of approximately 220 ± 70 CTX molecules *per* EV (Figure 2B). This was quantified by measuring the fluorescence of the Cyanine7.5 fluorophore, conjugated to CTX on RBC-EVs following SDS-PAGE analysis (Figure S2), and by correlating it with EV particle numbers determined by NTA. We obtained ^CTX^RBC-EVs with a mean diameter of 181 nm ± 6.7 nm, with no significant differences in size compared to the control (Figure S1D). To test EGFR-targeting ability of ^CTX^RBC-EVs, we first confirmed by flow cytometry the over-expression of EGFR in MDA-MB-231 TNBC cell line as compared to BT-474 cells as negative control (NC) (p= 0.0009, Figure 2C). Then, both pristine RBC-EVs and ^CTX^RBC-EVs were labeled with Memglow488, a fluorogenic, non-toxic membrane probe that fluoresces exclusively upon integration into biological membranes (27,53). MDA-MB-231 cells were incubated for 2 h at 4°C with two different amounts of RBC-EVs (i.e. 1✕10^9^ EVs/mL and 1✕10^10^ EVs/mL). (27) RBC-EVs at both tested concentrations were able to recognize whole MDA-MB-231 cells, as evidenced by the ≈100% of recognition displayed in Figure 2D. However, when we considered the values of mean fluorescence intensity (MFI), we evidenced a specific and dose-dependent contribution of CTX functionalization on cell recognition ability (Figure 2E). To date, it seems clear that a quote of non-specific binding occurred, since RBC-EVs can bind and recognize MDA-MB-231 cells even in absence of CTX functionalization. Considering that, we tried to define if the CTX on ^CTX^RBC-EVs was fully engaged in EGFR recognition by performing a binding assay, tracking the Cy7.5-labeled CTX, as previously described. As a control, free CTX was used in the same amount. Flow cytometry results evidenced that CTX on ^CTX^RBC-EVs was able to mark only a limited percentage of cells, despite in a dose-dependent manner (Figure 2F). Moreover, at the highest concentration, the CTX on ^CTX^RBC-EVs was more efficient in MDA-MB-231 binding than free CTX (Figure 2G). To define the specificity of this CTX-mediated binding, we performed a competition assay using free unlabeled CTX as a competitor. Flow cytometry evidenced that, in the fraction of ^CTX^RBC-EVs bound to MDA-MB-231 cells, there was almost a complete binding depletion in the presence of 100-fold molar excess of CTX, suggesting that the interaction between ^CTX^RBC-EVs and the cells was specifically mediated by EGFR (Figure 2H). To assess RBC-EV uptake by MDA-MB-231 cells, both pristine RBC-EVs and ^CTX^RBC-EVs were labeled with Memglow488. Given Memglow488 rapid diffusion into biological membranes, we also assessed RBC-EV uptake by analyzing the intracellular fluorescence signal resulting from the hemoglobin delivered by RBC-EVs. MDA-MB-231 cells were treated with Memglow488-labeled RBC-EVs and Memglow488-labeled ^CTX^RBC-EVs for 4 and 24 h. In both cases, we noticed a distinct intracellular fluorescence distribution concerning both Memglow488 and hemoglobin, with the former diffused throughout the cytoplasm, while the latter visible as distinct dots. These data are consistent with our previous work (27). (Figure 2I).

**Figure 2.**
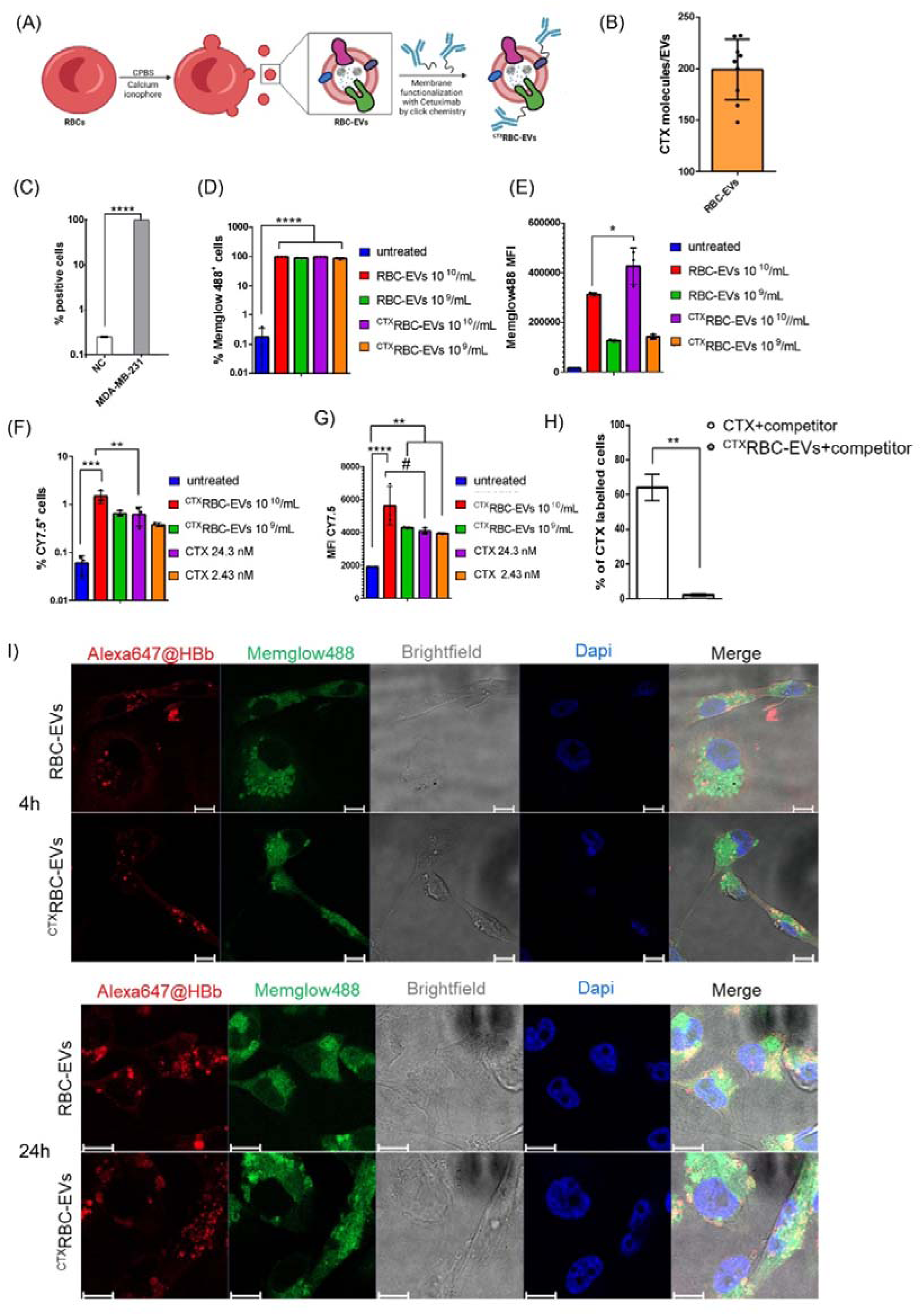
Characterization and *in vitro* uptake experiments of ^CTX^RBC-EVs on MDA-MB-231 TNBC cells. **(A)** Sketched scheme of the workflow followed for the production of ^CTX^RBC-EVs *via* bio-orthogonal click-chemistry. **(B)** CTX amount determination on the surface of RBC-EVs after click chemistry reaction. **(C)** Analysis of EGFR surface expression in MDA-MB-231 cell line by flow cytometry. BT474 cells were used as negative control (NC). ****, p= 0.0009**. (D and E)** Binding assay performed on MDA-MB-231 cell line with RBC-EVs and ^CTX^RBC-EVs labelled with Memglow 488. Results are expressed as percentage of Memglow 488 positive cells (D, ****, p= 0.0001) and MFI (E, *, p= 0.0109). **(F and G)** Binding assay performed on MDA-MB-231 cell line with Cy7.5 CTX free or immobilized on ^CTX^RBC-EVs. Results are expressed as percentage of Cy7.5 positive cells (F, ***, p= 0.0001 **, p=0.0067) and MFI (G, ****, p= <0.0001, **, p=0.0017, *, p=0.0331). **(H)** Competition assay performed with Cy7.5 CTX and ^CTX^RBC-EVs using a 100-fold molar excess of CTX as competitor. **(I)** Representative confocal microscopy images of MDA-MB-231 cells treated for 4 and 24 h with Memglow488-labeled RBC-EVs and Memglow488-labeled ^CTX^RBC-EVs. HBb, Hemoglobin, β subunit, scale bar: 10µm.

### Cisplatin encapsulates in ^CTX^RBC-EVs with the same encapsulation efficiency of RBC-EVs

After the optimization of cisplatin encapsulation and CTX functionalization procedures, we undertook the preparation and characterization of cisplatin-loaded ^CTX^RBC-EVs to produce the ^CTX^RBC-EV_cispt_ nanoplatform (Figure 3A). We first compared cisplatin encapsulation efficiency between ^CTX^RBC-EVs and RBC-EVs and we found no significant differences, leading to the conclusion that CTX functionalization did not hamper cisplatin encapsulation (Figure 3B). Then, we compared the size distribution of ^CTX^RBC-EV_cispt_ and ^CTX^RBC-EVs by NTA and we observed the presence of bigger NPs in ^CTX^RBC-EV_cispt_ (Figure 3C), as already described for RBC-EV_cispt_ (Figure 1D), with a mean diameter of 230 ± 12.2 nm for ^CTX^RBC-EV_cispt_ (27). Specifically, we observed statistically significant differences in size distribution between ^CTX^RBC-EVs, RBC-EVs, and ^CTX^RBC-EV_cispt_. (Figure S1D and Figure S3).

**Figure 3.**
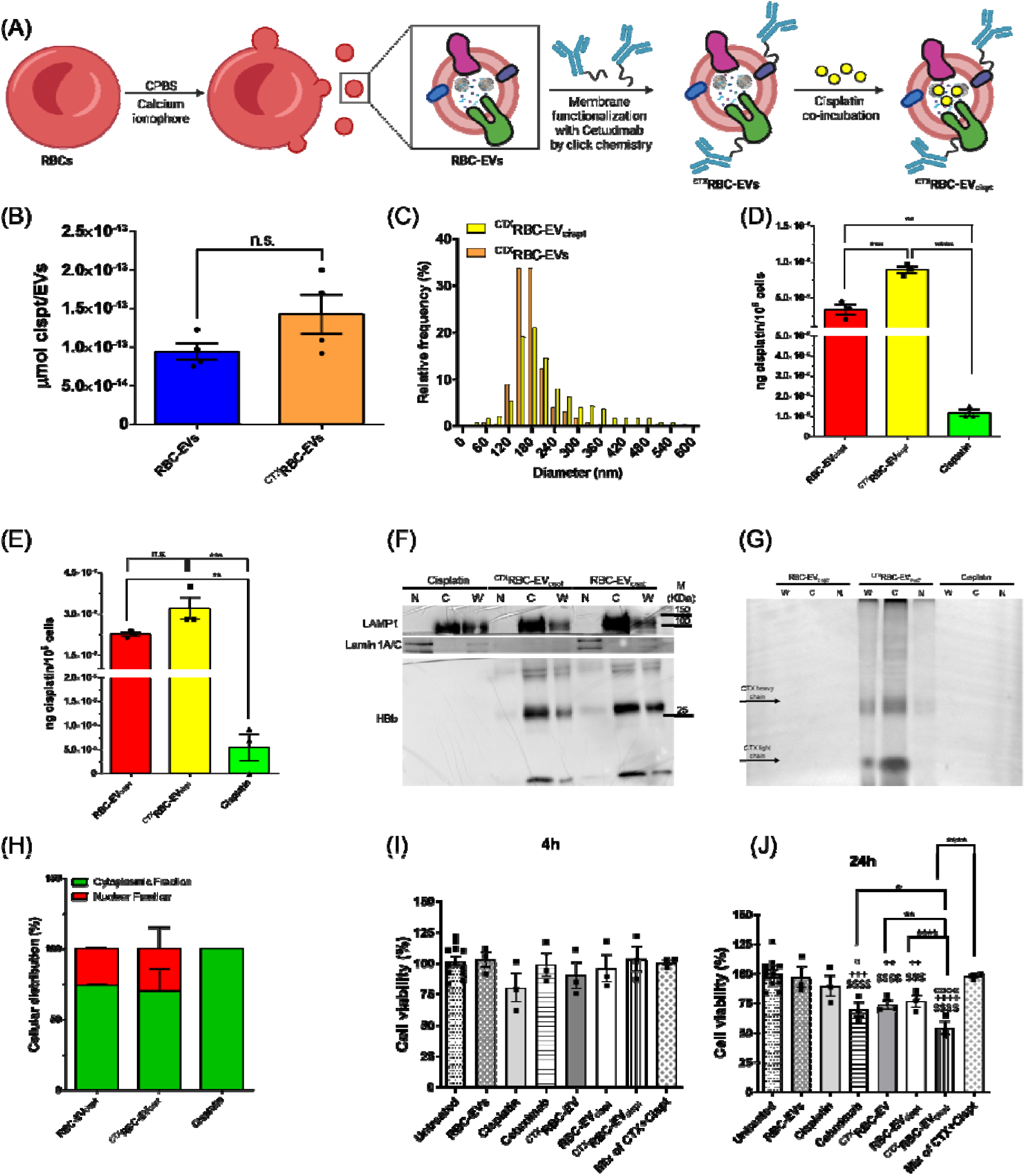
Characterization and *in vitro* uptake analysis of ^CTX^RBC-EV_cispt_ in MDA-MB-231 TNBC cells. **(A)** Schematics of functionalization and engineering process towards the production of ^CTX^RBC-EV_cispt_. **(B)** Comparison of cisplatin encapsulation yield between ^CTX^RBC-EVs and RBC-EVs after 24 h incubation. Unpaired t-test: *p* < 0.05, n.s. = not significant. **(C)** Size distribution of ^CTX^RBC-EV_cispt_ and ^CTX^RBC-EVs obtained by NTA measurements. **(D)** Quantification of intracellular cisplatin delivery after 30 min treatments with RBC-EV_cispt_, ^CTX^RBC-EV_cispt_ and free cisplatin (cisplatin final concentration = 5µM; Cetuximab final concentration = 20 nM). Ordinary one-way ANOVA test: p < 0.05, *****p < 0.0001. (**E**) Quantification of intracellular cisplatin delivery after 24 h treatments with RBC-EV_cispt_, ^CTX^RBC-EV_cispt_ and free cisplatin (cisplatin final concentration = 5µM; Cetuximab final concentration = 20 nM). Ordinary one-way ANOVA test: p < 0.05, *****p < 0.0001. **(F)** Western blot analysis of MDA-MB-231 cytoplasmic and nuclear fractions after 24h treatments with RBC-EV_cispt_, ^CTX^RBC-EV_cispt_ and free cisplatin. N = Nuclei, C = Cytoplasm, W = Whole cell lysate. (**G**) SDS-PAGE and fluorescence detection of Cy7.5-CTX in cytoplasmic and nuclear fractions obtained as in (F). (**H**) % of Cisplatin subcellular distribution in cytoplasmic and nuclear fractions obtained by dividing the amount of cisplatin in each fraction (ng) to the sum of cisplatin content in nuclear + cytoplasmic fractions (ng). **(I)** Cell viability analysis after 4h treatments. Data were normalized on untreated cells. Ordinary one-way ANOVA test showed no statistical significance. **(J)** Cell viability analysis after 24 h treatments. Ordinary one-way ANOVA test: $ = significance *vs*. untreated, + = significance *vs.* RBC-EVs, ° = significance *vs.* cisplatin, *p < 0.05, **p < 0.01, ****p < 0.0001.

### Cytotoxicity assessment of raw formulations: free cisplatin, pristine RBC-EVs and free CTX

In the perspective of developing a combined delivery system, we first assessed the cytotoxicity of the raw formulations, namely the free cisplatin, the pristine RBC-EVs and the free CTX. At first, we wanted to determine the sub-lethal doses of each component that can activate the mechanisms without strongly impacting cell viability. For the free cisplatin, we observed that cell viability declined in a dose- and time-dependent manner. We determined that the half-maximal inhibitory concentration (IC_50_) of cisplatin in MDA-MB-231 cells was approximately 36.41 ± 1.28 µM after 24 h and decreased to 12.3 ± 1.28 µM at 48 h (Figure S4A), consistently with the literature (51,54). The observed decrease in cisplatin IC_50_ over time in MDA-MB-231 cells could reflect cisplatin cumulative damaging effects. The difference in IC_50_ over time also indicated that the initial exposure to cisplatin at 24 h might not be sufficient to cause damage in all cells. However, with a longer incubation period, damage becomes more severe, leading to a larger proportion of cells undergoing death. We then analyzed the relationship between cell viability and their metabolic activity, measuring the oxygen consumption rate (OCR) with the Extracellular Flux Analysis technique in cells treated with increasing cisplatin concentrations. We observed that only the 25 µM cisplatin concentration displaying a toxic effect in MDA-MB-231 cells at 24 h (Figure S4A), significantly increased OCR values related to basal respiration, maximal respiration and spare respiratory capacity after 24 h in these cells (Figure S4B). This is consistent with a boosted energy supply and the activation of compensatory mechanisms aimed to counteract cisplatin-induce cytotoxicity. Although EVs are generally considered highly biocompatible, RBC-EVs have shown some inherent toxicity *per se* due to the high heme content in their lumen (55). To exclude any toxic effects deriving from RBC-EVs, we carried out a careful assessment of the appropriate RBC-EV doses and exposure times by MTT assay (Figure S4C). We observed a slight but not significant decrease in cellular viability only when cells were treated for 48 h with the highest concentration of RBC-EVs (10^11^ RBC-EVs/mL). Therefore, we decided to treat the cells for up to 24 h using an EV concentration of 6.7*10^10^ +/-2.21*10^10^ RBC-EVs/mL. Finally, we checked the effect of soluble CTX and obtained a slight but not significant decrease in cell viability for all tested concentrations (Figure S4D), which aligns with the literature (56). In line with cells viability data, evaluation of OCR showed that 24 h exposure to CTX did not cause any effect on mitochondrial respiration up to 800 nM. (Figure S4E). Considering both cell viability and metabolic data, we decided to use cisplatin and CTX at the final concentrations of 5µM and 10 nM – 40 nM, respectively, considering that - especially for the cisplatin - these doses were still able to induce cellular response without affecting cell viability.

### ^CTX^RBC-EV_cispt_ are uptaken more efficiently than RBC-EV_cispt_ in MDA-MB-231 cell line

Firstly, we checked the cellular uptake of both RBC-EV_cispt_ and ^CTX^RBC-EV_cispt_ by immunofluorescence analysis at 4 and 24 h. The results were consistent with those obtained for ^CTX^RBC-EVs and RBC-EVs, thus confirming that ^CTX^RBC-EV_cispt_ were taken up by cells already after 4h (Figure S5A) and hemoglobin signal persisted up to 24h (Figure S5B) (27). We next compared the amount of cisplatin delivered into cells by RBC-EV_cispt_ and ^CTX^RBC-EV_cispt_ or in its free form by ICP-MS. This powerful technique allows us to precisely quantify the amount of cisplatin delivered into cells without the need to modify its structure by attaching fluorophores. We incubated MDA-MB-231 cells with the three preparations at a sublethal dose of cisplatin (5 µM) for 30 minutes and 24 h. The change in uptake time points (from 4 h to 30 minutes) was experimentally motivated by the observation that, at 4 h, uptake signals were very similar between RBC-EVs and ^CTX^RBC-EVs. Therefore, we decided to assess cisplatin uptake at 30 minutes to confirm targeted uptake at an earlier time point. After 30 minutes, cisplatin was already detected in cells, especially delivered *via* RBC-EVs (Figure 3D). A statistically significant difference was observed between RBC-EV_cispt_ and ^CTX^RBC-EV_cispt_, with a 3-fold in cisplatin delivery trough ^CTX^RBC-EV_cispt_ compared to RBC-EV_cispt_. This suggests that ^CTX^RBC-EV_cispt_ may utilize a more efficient or targeted uptake mechanism, likely facilitated by the interaction of CTX with its EGFR on the cell surface (Figure 2). After 24 h, the amount of intracellular cisplatin was still detectable and higher when it was delivered by RBC-EV_cispt_ or ^CTX^RBC-EV_cispt_ compared to the free drug. However, we did not observe significant differences between RBC-EV_cispt_ and ^CTX^RBC-EV_cispt_ at this time point (Figure 3E), suggesting that the initial advantage of ^CTX^RBC-EV_cispt_ in rapid uptake does not necessarily translate into long-term differences in intracellular cisplatin concentration. Taken together, these findings underscore a remarkable enhancement in intracellular cisplatin delivery, with RBC-EV_cispt_ and ^CTX^RBC-EV_cispt_ achieving an approximately 40- to 60-fold increase compared to the free drug and ^CTX^RBC-EV_cispt_ showing a more efficient initial uptake, possibly due to targeted endocytosis.

### Cisplatin and cetuximab are delivered into cells by ^CTX^RBC-EV_cispt_ nanoformulation

Since cisplatin can exert toxicity at both nuclear and cytoplasmic level, we checked its subcellular distribution in MDA-MB-231 cells. Cells were treated for 24 h either with free cisplatin or RBC-EV_cispt_ and ^CTX^RBC-EV_cispt_ and then their nuclear fraction was separated from the cytoplasm, as explained in the methods section (33). Western blot analysis confirmed the subcellular fractionation, highlighting the presence of the Lysosomal Associated Membrane Protein 1 (LAMP1) exclusively in the whole cellular homogenate and in the cytoplasmic fraction, while lamin A/C (a protein expressed in the nuclear lamina) was detected, as expected, in the whole homogenate and in the nuclear fraction (Figure 3F). Hemoglobin (HBb) was found mostly expressed in the cytoplasm (confirming the imaging data), although a low signal was also detected in the nuclear fraction (Figure 3F). This suggests that some RBC-EV-derived components might translocate into the nucleus, although the mechanism and significance of this minor nuclear presence remain unclear. Interestingly, we noticed that cells treated with ^CTX^RBC-EV_cispt_ showed a low expression of lamin A/C. The lamin A/C reduction could indicate changes in nuclear structure or integrity, potentially influencing nuclear functions such as DNA repair and cell cycle regulation (57). This could be a direct result of the targeted delivery mechanism of ^CTX^RBC-EV_cispt_, which may facilitate a more pronounced impact also on nuclear pathways as compared to free cisplatin or RBC-EV_cispt_. We also checked the intracellular localization of CTX by analyzing the fluorescence signal of the Cy7.5 bound to CTX. We observed that, although the majority of CTX was located in the cytoplasm, a small amount was also detected in the nuclear fraction (57) (Figure 3G). We then quantified the nuclear/cytoplasmic distribution of cisplatin by ICP-MS and normalized the amount of cisplatin in each fraction on the total amount of cisplatin detected in that sample (cytosolic cisplatin+ nuclear cisplatin) (Figure 3H). In cells treated with the free drug, 100% of the total cisplatin was localized in the cytoplasm, with no cisplatin detected in the nuclei. In cells treated with RBC-EV_cispt_ or ^CTX^RBC-EV_cispt_, still most of the cisplatin was found in the cytoplasm (≈ 75% and ≈ 70%, respectively), although a nuclear localization is also present, indicating that the different route of cisplatin administration changes its subcellular localization. This suggests that cisplatin primary mode of action could still be largely cytoplasmic, particularly at the sub-lethal concentration of 5 µM. Furthermore, the localization of cisplatin in the nucleus of cells, together with hemoglobin and cetuximab, suggests the endocytosed RBC-EVs can deliver their cargo to the nucleus, as previously reported (58).

### MDA-MB-231 viability is significantly decreased upon treatment with ^CTX^RBC-EV_cispt_

We assessed the potential of the ^CTX^RBC-EV_cispt_ nanoformulation as anti-proliferative system on MDA-MB-231 cells for 4h a 24h using the doses previously determined (final cisplatin concentration 5 µM, using 6.71 *10^10^ prt/ml +/- 2.21 *10^10^ prt/ml, Figure S4). 4h and 24h time points were chosen to better assess the metabolic effects of the formulations, which might not be observed at earlier time points. After 4h, both RBC-EV_cispt_ and ^CTX^RBC-EV_cispt_ demonstrated a slight effect on cellular viability as compared to the free cisplatin or CTX, without statistically significant differences (Figure 3I). After 24h, both RBC-EV_cispt_ and ^CTX^RBC-EV_cispt_ significantly decreased cell viability as compared to the untreated cells and the cells treated with pristine RBC-EVs. However, treatments carried out with ^CTX^RBC-EV_cispt_ showed the most critical impact, resulting in a reduction of approximately 50% of cell viability, differently from cells treated with RBC-EV_cispt_ (Figure 3J). We also measured OCR in cells treated with RBC-EV_cispt_ and ^CTX^RBC-EV_cispt_, and we observed that none of the two conditions affected OCR in MDA-MB-231 cells (Figure S4F). No change in Extracellular Acidification Rate (ECAR) was furthermore detected within all the experiments tested (data available upon request).

### ^CTX^RBC-EV_cispt_ sensitize cancer cells to oxidative stress and promote ferroptosis

To investigate the molecular mechanisms underlying the observed cytotoxic and cellular effects of the different formulations, we performed the RNA-seq transcriptome profiling of MDA-MB-231 cells subjected to the different treatments for 24 h (namely, RBC-EVs, RBC-EV_cispt_, ^CTX^RBC-EVs, ^CTX^RBC-EV_cispt_, free cisplatin, free CTX, and a mix of free CTX + free cisplatin) as well as of untreated cells (wild type) as control. From the sequencing, we obtained at least 21 millions (M) read pairs/sample, with a mean of 94% uniquely mapped reads on genome and 84% reads uniquely assigned to genes (Suppl File 1). Concerning the single-agent treatments, differential expression analysis did not find any differentially expressed genes (DEGs, adjusted p-value < 0.1 and |log2FC| >1) in cells treated with RBC-EVs as compared to untreated cells, and only one DEG in free CTX-treated *vs.* untreated cells, confirming the non-toxicity of these two agents at such doses and times. On the other hand, free cisplatin-treated cells showed 11 DEGs as compared to untreated cells (Table 1 and full lists in Suppl File 2). For the combined treatments, we identified 22 DEGs for the mix of free CTX + free cisplatin, 22 DEGs for RBC-EV_cispt_ and 19 DEGs for ^CTX^RBC-EV_cispt_ as compared to untreated cells (Table 1 and full lists in Suppl File 2). Interestingly, three DEGs were deregulated by both cisplatin and the mix of CTX and cisplatin treatments (PARD6G as up- and SHANK2 and RPLP0P9 as down-regulated DEGs), and two DEGs were in common between RBC-EV_cispt_ and ^CTX^RBC-EV_cispt_ treatments (CYP1B1 as up- and STC1 as down-regulated DEGs).

**Table 1.**
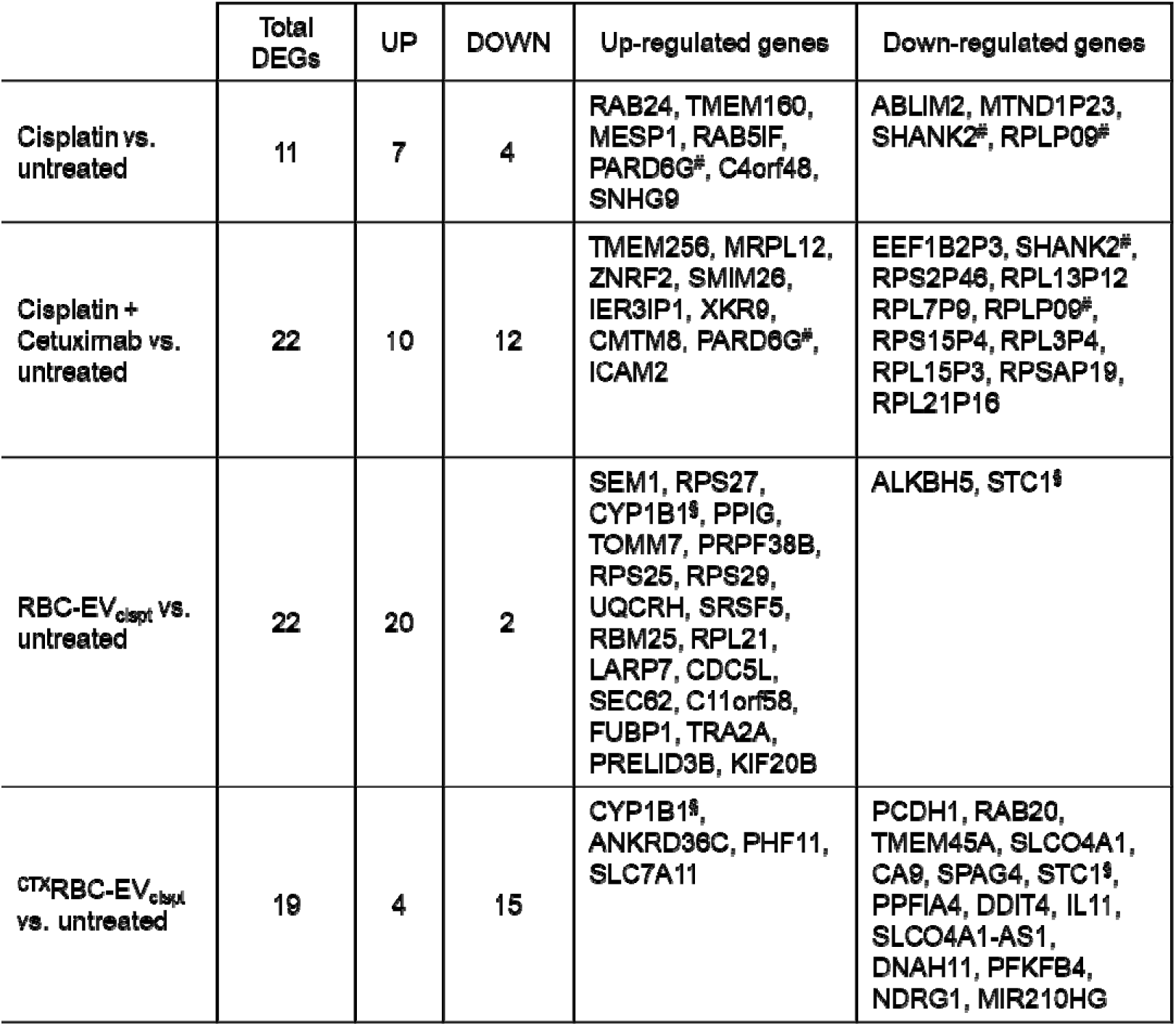
DEG counts (adjusted p-value < 0.1 and |log2FC| > 1). For each comparison, DEGs are listed according to decreasing fold-change values. RBC-EVs, red blood cell-derived extracellular vesicles; wt, untreated cells; CTX, Cetuximab; cispt, Cisplatin; ^#^, common genes between Cisplatin and the mix of CTX and cisplatin treatments; ^§^, common genes between RBC-EV_cispt_ and ^CTX^RBC-EV_cispt_ treatments.

To explore the functional processes altered by the different formulations and related to the cytotoxic effects in MDA-MB-231 treated cells, we performed the functional enrichment analysis of DEGs on Gene Ontology Biological Process (GO-BP) terms and several pathway collections (Suppl File 3).

Interestingly, we found that cisplatin, administered as both free drug and mix of CTX and cisplatin, increased apical junction interactions as well as signaling pathways mediated by Hippo and Notch (Figure 4A, and full list of enrichments in Suppl File 3). In particular, after treatment with free cisplatin, we found the up-regulation of Mesoderm Posterior BHLH Transcription Factor 1 (MESP1) and Partitioning Defective 6 Homolog Gamma (PARD6G) genes (Suppl File 3). MESP1 is a transcription factor with a crucial role in epithelial–mesenchymal transition (EMT), Notch signaling and cell migration, and EMT has been linked to the activation of HIF-1α signaling, which can enhance cell motility, support cellular nutrient metabolism, and stimulate angiogenesis to boost energy supply, enabling cancer cells to survive in challenging conditions (59). On the other hand, PARD6G is a key component of the Par6/aPKC complex, which regulates cell polarity, apical junction formation, asymmetric cell division and Hippo signaling, and it is correlated to cancer development (60). As for MESP1, PARD6G up-regulation may have several implications in the context of TNBC and cisplatin response. For instance, it suggests the activation of compensatory mechanisms, where cells attempt to restore post-EMT polarity and adhesion, potentially enhancing motility or invasiveness. All these functions are already known mechanisms associated with an increased cisplatin resistance in cancer (61).

**Figure 4.**
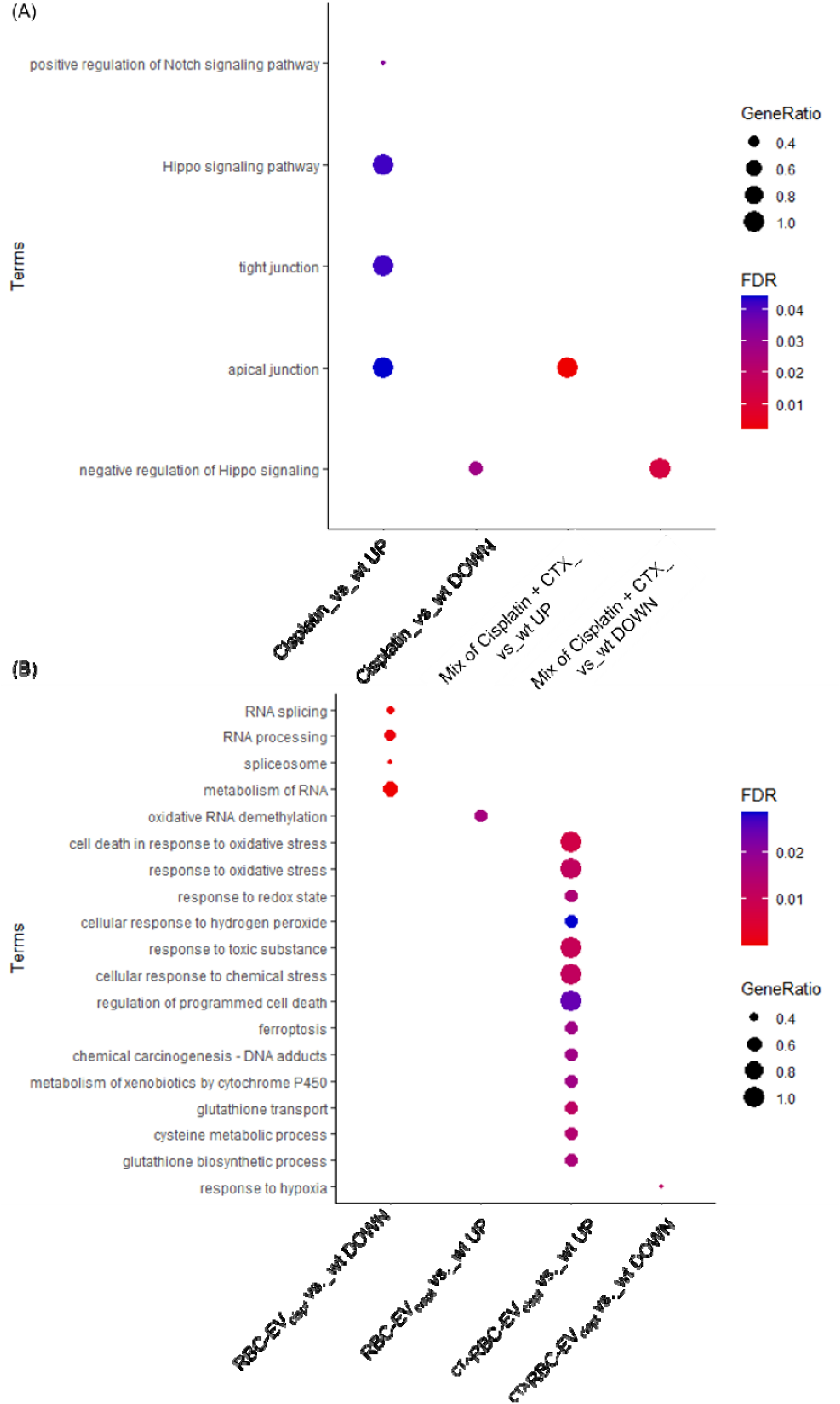
Functional enrichment analysis in MDA-MB-231 cells treated for 24h with the different formulations. Bubble plots show a selection of biological functions of interest significantly enriched by up- and down-regulated genes in cells treated with **(A)** free cisplatin and mix of free cisplatin + free CTX (Mix of Cisplatin and CTX), and **(B)** RBC-EV_cispt_ and ^CTX^RBC-EV_cispt_. Bubble size represents the gene ratio for each enriched function, while color gradient indicates statistical significance of the enrichment (FDR-BH).

We also observed the down-regulation of SH3 And Multiple Ankyrin Repeat Domains 2 (SHANK2) gene, involved in the negative regulation of the Hippo signaling pathway (Figure 4A). The Hippo pathway plays a crucial role in regulating cell proliferation, apoptosis, and ferroptosis through its effectors YAP and TAZ (62) . SHANK2 down-regulation alters this pathway, potentially diminishing the effectiveness of ferroptosis-based therapies (63). Furthermore, in TNBC, Hippo pathway is shown to induce cisplatin resistance *via* EMT activation. Thus, FDA-approved drugs that indirectly block Hippo/YAP signaling are showing promise in the clinic for reducing cancer chemoresistance (64).

We found PARD6G up-regulation and SHANK2 down-regulation also after treatment with the mix of free cisplatin and free CTX, in addition to the modulation of other interesting genes, such as the mitochondrial ribosomal protein L12 (MRPL12) gene, highlighting alterations in mitochondrial functions and possibly energy metabolism, which are critical in maintaining cell survival under stress conditions like chemotherapy. Finally, no term related to drug cytotoxic effects was found enriched when cisplatin was administered as a free drug (as both free cisplatin and a mix of CTX and cisplatin), thus supporting our observations on cell viability after free drug treatments described above.

Differently, when MDA-MB-231 cells were treated with RBC-EV_cispt_ and, even more, with ^CTX^RBC-EV_cispt_, we observed many enriched terms concerning drug cytotoxic effects (Figure 4B and full list of enrichments in Suppl File 3). Indeed, cells treated with RBC-EV_cispt_ activated a group of functions related to RNA splicing, RNA processing, spliceosome and RNA metabolism (with up-regulation of PPIG, PRPF38B, SRSF5, RBM25, LARP7, CDC5L, TRA2A, RPS27, RPS25, RPS29, SEM1, RPL21), which have been functionally linked to the cell death induced by cisplatin (65) (Figure 4B and Figure S6A). In particular, SEM1 and Ribosomal Protein S27 (RPS27) genes are both involved in protein degradation and ribosome function, suggesting an increased protein synthesis and cellular stress response, which is crucial when cancer cells are exposed to cisplatin. Other notable up-regulated genes include TOMM7, involved in mitochondrial function and protein import into the mitochondria, thus suggesting alterations in mitochondrial activity, critical in cellular stress response during chemotherapy (66). Interestingly, we found the down-regulation of oxidative RNA demethylation function (Figure 4B), involving ALKBH5 gene, a known RNA demethylase contributing to RNA stability and methylation, processes critical for gene expression regulation. ALKBH5 down-regulation might affect cancer cell proliferation and survival, potentially increasing their sensitivity to ferroptosis and contributing to the effectiveness of cisplatin-loaded RBC-EVs (67).

Even more, when treated with ^CTX^RBC-EV_cispt_, MDA-MB-231 cells showed increased oxidative stress, formation of DNA adducts and cell death (Figure 4B), thus confirming the higher cytotoxicity of cisplatin vehiculated in this manner, resulting from an enhanced uptake and delivery into cells as well as trafficking towards the nuclei as above described. Nonetheless, cells treated with ^CTX^RBC-EV_cispt_ showed also the up-regulation of terms related to response to toxic substance and chemical stress and to metabolism of xenobiotics by cytochrome P450 (Figure 4B), thus indicating the activation of detoxification mechanisms to counteract cisplatin activity.

Finally, treatment with ^CTX^RBC-EV_cispt_ showed to affect several processes related to an enhanced ferroptosis and a decreased hypoxia response (Figure 4B), two functions that contribute to reduce chemoresistance and to sensitizing cancer cells to cisplatin. In particular, we found the up-regulation of CYP1B1 gene, a member of the cytochrome P450 family involved in xenobiotic metabolism and previously associated with chemotherapy resistance in cancer cells (68) (Figure S6B). Interestingly, CYP1B1 is also involved in the metabolism of polyunsaturated fatty acids, leading to the production of reactive oxygen species (ROS) that can promote lipid peroxidation and ferroptosis. We also found the up-regulation of SLC7A11 (Figure S6B), which is essential for the uptake of cystine, a precursor for the antioxidant glutathione (GSH). When SLC7A11 is up-regulated, there is an increase in cystine uptake, leading to elevated GSH intracellular levels, which helps in maintaining cellular redox homeostasis (as illustrated in KEGG ferroptosis map 04216 (10,69) (Figure S7). Conversely, if cisplatin depletes GSH, it can create a feedback loop where the expression of SLC7A11 might be further increased in the attempt to restore GSH levels.

Interestingly, we also found a slight down-regulation of EGFR, that might lead to sensitize cells to chemotherapy and potentially contribute to HIF-1a regulation and ferroptosis activation. In fact, the down-regulation of the Signal Transducer and Activator of Transcription 3 (STAT3) is crucial in EGFR-HIF1a cross-talk. STAT3 is a survival and oncogenic factor frequently activated downstream of EGFR (as illustrated in KEGG EGFR map 01521 (Figure S8) and its down-regulation suggests a disruption in pro-survival signaling cascades. STAT3 is known to promote cell proliferation, inhibit apoptosis, and regulate HIF-1a under hypoxic conditions (as illustrated in KEGG HIF-1 map 04066 (Figure S9). Its down-regulation might interfere with HIF-1a stabilization, decreasing the expression of genes associated with cell survival, hypoxia response and resistance to oxidative stress, thereby sensitizing cells to ferroptosis. In this context, we coherently found the down-regulation of Carbonic anhydrase IX (CA9), Stanniocalcin 1 (STC1), DNA Damage Inducible Transcript 4 (DDIT4), N-myc Downstream Regulated Gene 1 (NDRG1), PTPRF Interacting Protein Alpha 4 (PPFIA4) and Phosphofructo-Kinase/Fructose-Biphosphatase 4 (PFKFB4) genes, which are all regulated by HIF-1a in hypoxia response (as showed by protein-protein interaction network by STRING (Figure 5) and Cytoscape (Figure S10). Among them, NDRG1 gene is implicated in cellular stress responses, including iron homeostasis, hypoxia response and oxidative stress management and its down-regulation may enhance susceptibility to ferroptosis, while DDIT4 is involved in autophagy and stress responses (70), and its down-regulation can reduce autophagic flux and increase sensitivity to ferroptosis.

**Figure 5.**
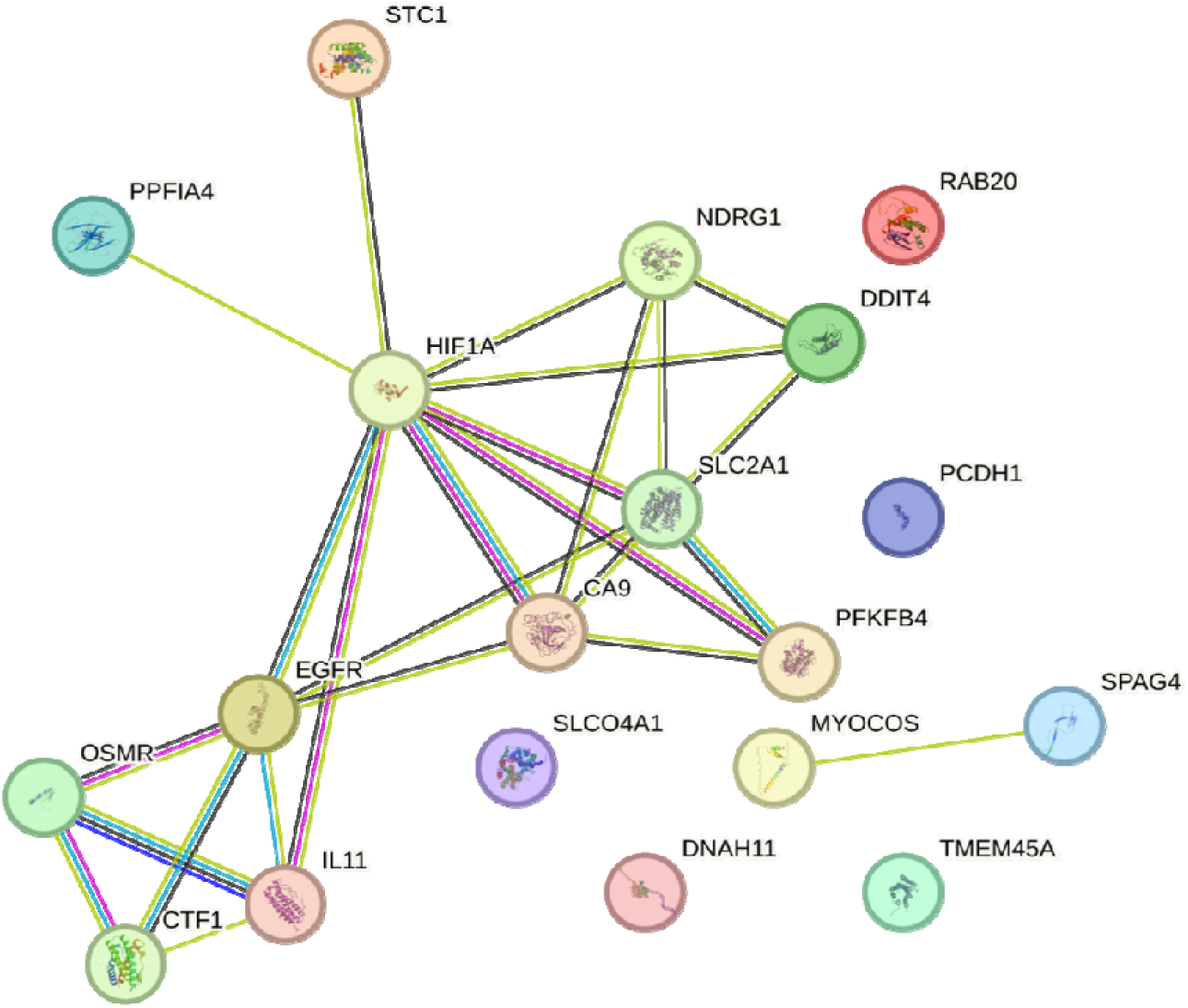
STRING network of protein-protein interactions among the down-regulated genes in ^CTX^RBC-EV_cispt_ *vs.* wt cells. The network highlights the genes controlled by HIF-1a in hypoxia response cascade. Interactions are colored according to STRING default settings (v.12.0).

In our opinion, the observed down-regulation of EGFR, STAT3, CA9, STC1, DDIT4, NDRG1, PPIA4, and PFKFB4 genes in TNBC cells treated with ^CTX^RBC-EV_cispt_ may reflect the disruption of key survival and stress response pathways. This may impair HIF1a-mediated hypoxic adaptation, thus sensitizing cancer cells to oxidative stress and promoting ferroptosis.

Based on these observations, we assessed ferroptosis promotion based on lipid peroxidation in the MDA-MB-231 cell line performing the following treatment with: RBC-EVs, RBC-EV_cispt_, CTX-RBC-EVs, ^CTX^RBC-EV_cispt_, free cisplatin, free CTX, and a combination of free CTX and free cisplatin. Untreated cells were used as a negative control, while Erastin-treated cells served as a positive control (71,72). Lipid peroxidation levels were evaluated using C11-BODIPY^581/591^: upon oxidation, the fluorescence of this fluorophore shifts from red to green (40,43). The green-to-red fluorescence intensity ratio was calculated as an oxidation index to quantify lipid peroxidation (Figure 6A and S12). Statistical analysis of the oxidation index, comparing all treatments to untreated cells, revealed that both the combination of free CTX and cisplatin (Mix of CTX + Cispl) and ^CTX^RBC-EV_cispt_ induced a significant increase in lipid peroxidation, indicative of ferroptosis, as shown in Figure 6B. None of the other treatments increased lipid peroxidation levels compared to the negative control, whereas the Erastin treatment caused a significant increase in peroxidase activity, confirming its effectiveness as a positive control. Taken together these results indicate that *in vitro* analyses support the findings from the RNAseq analysis: MDA-MB-231 cells treated with ^CTX^RBC-EV_cispt_ exhibit increased oxidative stress, suggesting the promotion of ferroptosis. Concerning the treatment with free CTX and cisplatin (Mix of CTX + Cispt) RNA seq data showed that gene SLC7A11 resulted up-regulated also in the Cispt+CTX vs wt comparison (log2FC = 0.6, padj = 2.12E-05), although it does not pass the thresholds imposed on the log2FC value, and therefore it was not classified as DEG. This might be explained by the fact that many core regulators of ferroptosis are modulated at the transcriptional (i.e. miRNAs), post-transcriptional, translational and post-translational levels (73–75), so a clear change in RNA-seq for that gene cannot be demonstrated in this case. Previous data only established cetuximab as a sensitizer to ferroptosis, but not specifically in combination with cisplatin (76). The only available data indicate that in head and neck cancer, the regimen cetuximab + cisplatin is shown to drive mainly apoptotic and immunogenic cell death but ferroptosis markers were not assessed (77).

**Figure 6.**
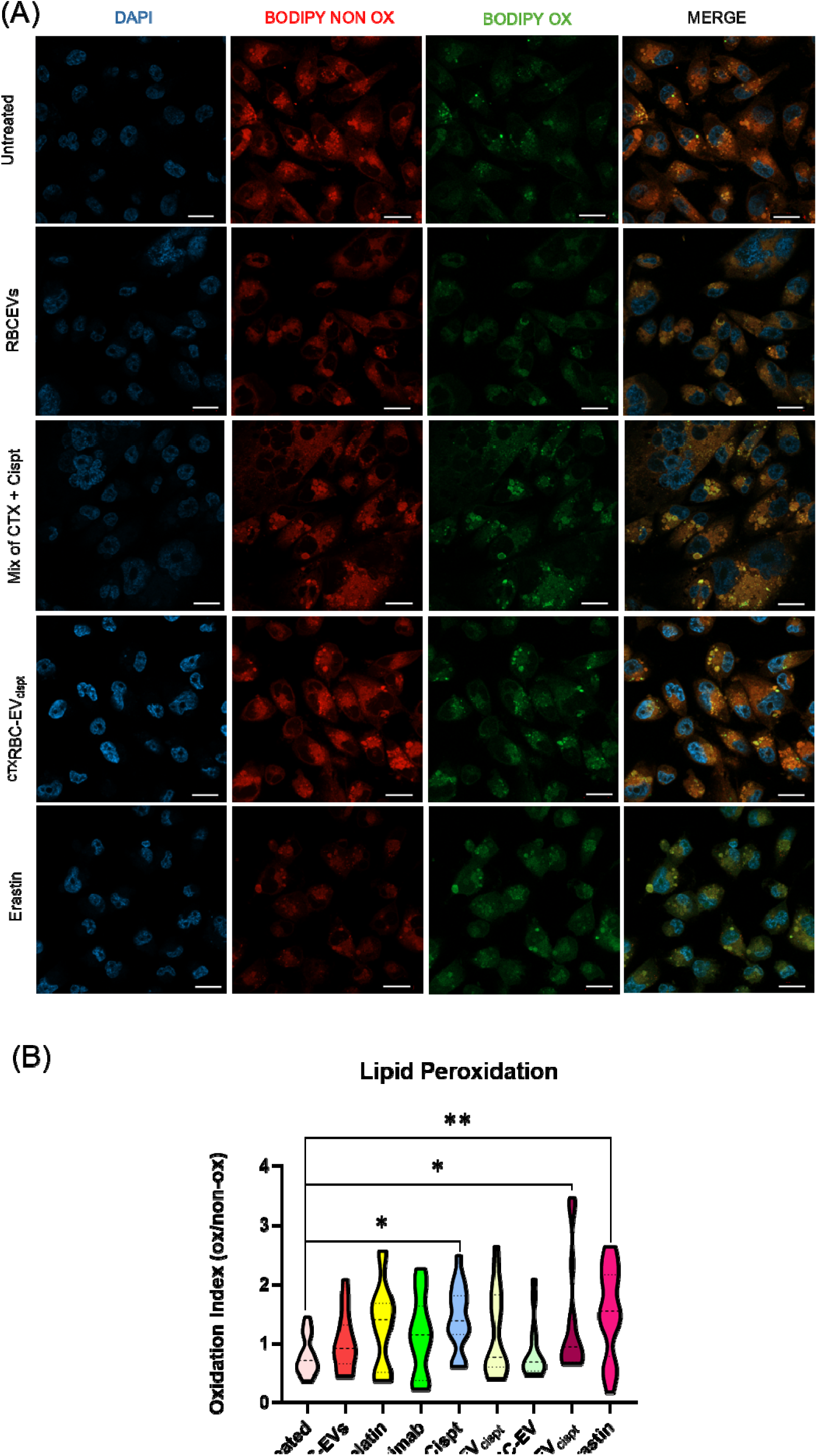
Ferroptosis promotion evaluation via lipid peroxidation. (A) Representative confocal microscopy images of cells stained with BODIPY™ 581/591 C11 (oxidized: green; non-oxidized: red) to visualize lipid peroxidation, and DAPI (blue) to label nuclei. Merged images show all channels combined. Treatments include untreated control (Untreated), RBC-EVs (vehicle), treatment with: free CTX and cisplatin (Mix of CTX + Cispt), ^CTX^RBC-EV_cispt_ , or Erastin as a positive control for ferroptosis induction. Images are shown for selected conditions. Scale bar 20 um. (B) Quantification of lipid peroxidation as oxidation index (ratio of oxidized [green] to non-oxidized [red] BODIPY fluorescence), displayed as violin plots representing the full distribution of values across all tested conditions and multiple fields of view. Statistical analysis was performed using ordinary one-way ANOVA followed by Dunnett’s multiple comparisons test vs untreated control. Statistically significant differences are indicated as: *p < 0.05, **p < 0.01;* unmarked conditions are not statistically significant.

### ^CTX^RBC-EV_cispt_ nanoplatform limited cisplatin *ex vivo* toxicity by limiting the production of ghost red blood cells

In addition to chemoresistance, off-target toxicity represents one of the major drawbacks of cancer therapies. In particular, cisplatin-based treatments suffer from numerous systemic toxicity effects, among which hemotoxicity and anemia are considered ones of the most diffused ones (78–81). For cisplatin, anemia can occur because of the ability of cisplatin to interact with red blood cell plasma membrane (59), causing (i) a reduction in the number of erythrocytes and (ii) morphological changes towards stomatocytic shape (81). We performed an *ex vivo* toxicity assay, by incubating red blood cells with ^CTX^RBC-EV_cispt_ (5 µM cisplatin) and soluble cisplatin at both sub-lethal (5 µM) and lethal doses (25 µM) for 3 h at 37°C. In all samples, we observed the presence of erythrocytes with altered morphology (Figure 7, yellow arrows, i.e., stomatocytes and acanthocytes), although with different percentages (17.1 ± 2.2 % for untreated, 23.2 ± 2.5% for the vehicle, 43.1 ± 4.3% for 5 µM cisplatin, 34.3 ± 12.7% for 25 µM cisplatin and 32.5 ± 0.1% for ^CTX^RBC-EV_cispt_). Ghost RBCs were found only after treatments with the free drug, reaching the highest percentages when erythrocytes were incubated with 25 µM cisplatin (Figure 7, white arrows). Altogether, these results demonstrated the low hemotoxicity of ^CTX^RBC-EV_cispt._ nanoplatform. These findings are consistent with, and further supported by, previous results from our laboratory and collaborators about RBC-EV safety. The immunotoxicity of RBC-EVs was assessed in THP-1 M(0) cells exposed to increasing concentrations of RBC-EVs for 24 and 48 h in complete RPMI-1640 medium. No cytotoxic effects were detected at either time point, confirming their safety profile (82). In addition, RBC-EVs were evaluated in vivo using a *C. elegans* model, which demonstrated not only the absence of toxicity but also a significant increase in locomotor activity after 72 h of treatment, likely attributable to the antioxidant enzymes present within RBC-Evs (83).

**Figure 7.**
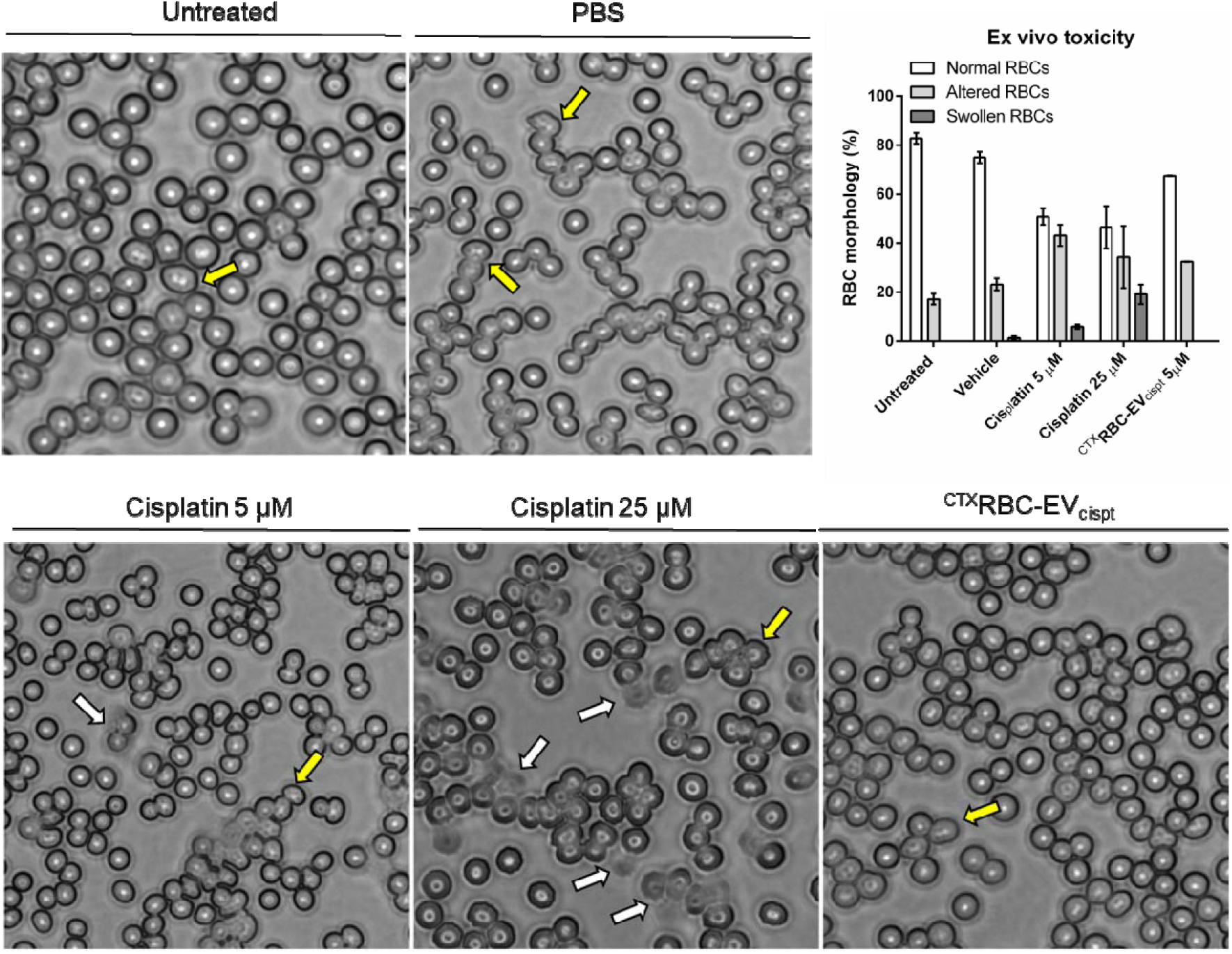
*Ex vivo* morphological analysis of red blood cells after 3 h treatments with cisplatin and ^CTX^RBC-EV_cispt_. Yellow arrow = altered RBCs, white arrow = ghost RBCs. Cell counting was performed with Fiji software and the relative percentage of RBC morphologic alterations were calculated as follow: (Number of RBCs with a specific morphology/Total counted RBCs) x 100. Data are expressed as mean ± SD.

### Action of ^CTX^RBC-EV_cispt_ in *ex vivo* 3D TNBC patient-derived organoids

To demonstrate the activity of engineered RBC-EVs as tumor targeted delivery vehicle of Cisplatin, we decided to assess their performances on two Patient-Derived Organoids (PDOs) models of TNBC, PDO#1 and PDO#2, which were fully characterized (Figure S11). PDO#1 and PDO#2 were both derived from invasive ductal carcinoma with a TNBC molecular subtype, despite only PDO#1 displayed BRCA1 mutation. We performed a cell viability assay to compare the efficacy between free Cisplatin and the cisplatin-based nanoformulations RBC-EV_cispt_ and ^CTX^RBC-EV_cispt_, using two different drug concentrations (5 µM and 25 µM). After 72 h of treatment, PDO#1 was sensitive to both doses of Cisplatin (p<0.0001), while nanoformulated Cisplatin seemed to be ineffective in this model, with the only exception of 25 µM RBC-EV_cispt_, which evidenced a low but statistically significant ATP reduction (Figure 8). Moreover, in this model the functionalization with CTX (^CTX^RBC-EV_cispt_) had no effect (Fig. 8), as expected considering the low expression of EGFR in PDO#1. In contrast, PDO#2 was only sensitive to high doses of Cisplatin, RBC-EV_cispt_ and ^CTX^RBC-EV_cispt_ (25 µM) (Figure 8). These findings indicate that, in the case of low sensitivity to Cisplatin, the functionalization with CTX results in higher treatment efficacy and better intracellular penetration. Therefore, we can assume that the multimodal engineering of RBC-EVs with Cisplatin and CTX promotes drug uptake within the organoid and, consequently, reduces cell viability.

**Figure 8.**
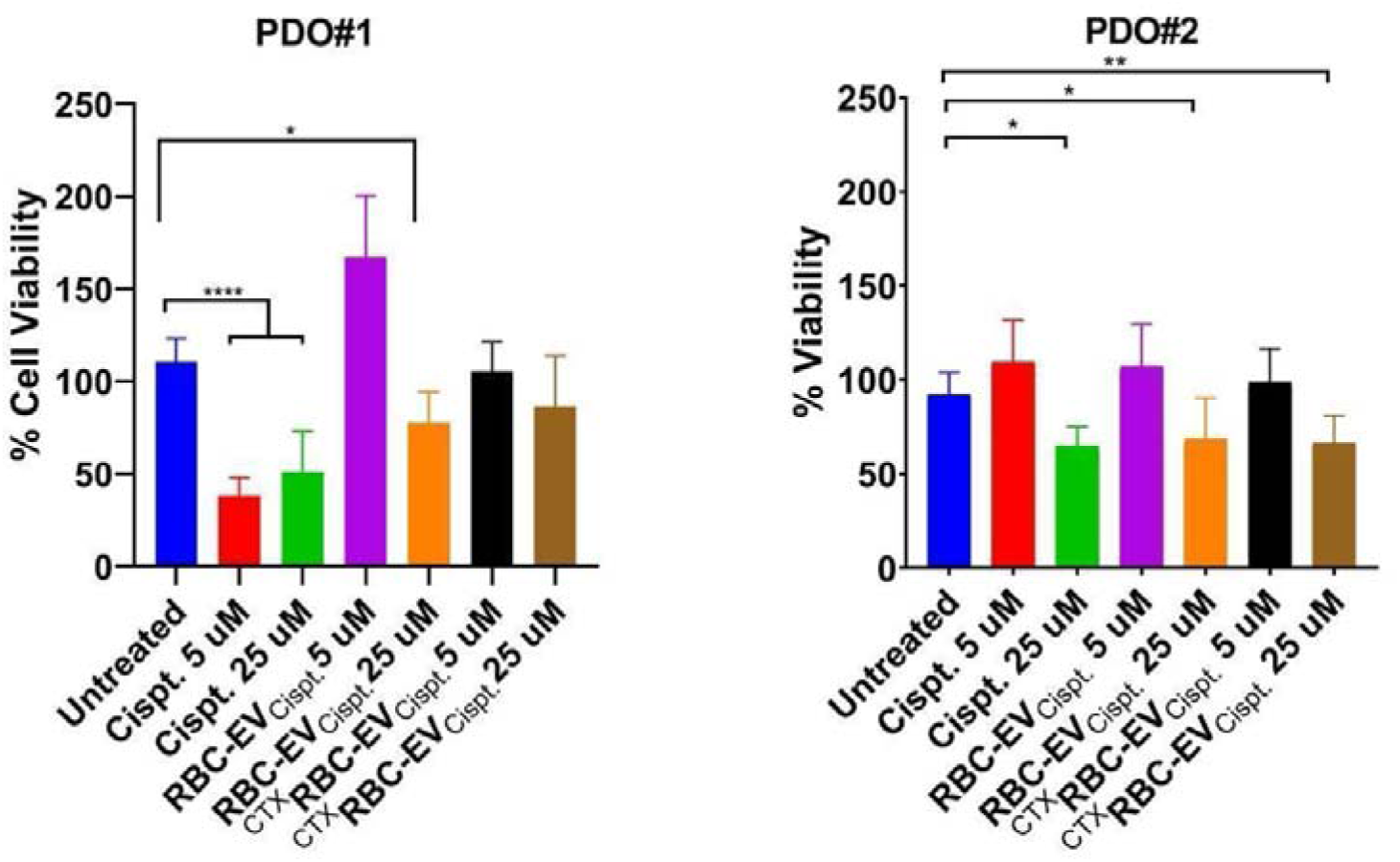
Cell Viability assay to assess sensitivity of PDO#1 and PDO#2 to Cisplatin (Cispt.), RBC-EV_Cispt_ and ^CTX^RBC-EV_Cispt_ at two different concentrations (5 and 25 μM). Ten replicates for each condition were tested, at least two biological replicates were performed. Untreated organoids were used as negative control. The data are reported as mean ± SD; *p<0.05, **p<0.001, ****p<0.0001.

## Conclusions

Despite its potential sensitivity to cisplatin, TNBC treatments are still challenged by cisplatin-acquired chemoresistance.

Nevertheless, the combination of RBC-EVs, Cetuximab and Cisplatin presents a promising therapeutic strategy for treating TNBC by inducing cell death through ferroptosis. Our RNA-seq data suggested a possible mechanism of action for our ^CTX^RBC-EV_cispt_ nanoplatform. Cetuximab, a monoclonal antibody targeting the epidermal growth factor receptor, which is highly over-expressed in our TNBC cancer cell model, blocks EGFR signaling cascade, leading to the down-regulation of hypoxia-inducible factor 1-alpha (HIF-1A) downstream genes. This reduction in HIF-1A downstream cascade impairs the tumor ability to adapt to hypoxic conditions, making cancer cells more vulnerable to oxidative stress. Concurrently, cisplatin can exert its cytotoxic effects by binding to and depleting glutathione (GSH), a critical antioxidant. The GSH depletion increases oxidative stress, which, combined with HIF-1A hypoxia response down-regulation, creates an environment conducive to ferroptosis and, consequently, to cell death. This multifaceted mechanism underscores the potential of combining RBC-EVs, cetuximab and cisplatin, thus enhancing their therapeutic efficacy against TNBC by promoting ferroptosis, ultimately leading to increased tumor cell death and preventing anemia. Further investigations on this drug combination may provide valuable insights for improving treatment efficacy in TNBC.

## Supporting information

RNA seq metrics

DEG list

Functional analysis

enrichment

## Supporting Information

The Supporting Information is available free of charge.

## Author information

Conceptualization: A.R., M.R. A.M, S.M., I.C., P.B; data curation: M.R., S.M., I.C., A.S. and A.R., Funding acquisition: A. R., P. B. and F.C.; Investigation: M.R., A. M., A. Z., L. P., S. A., R.Z., S. T., I.C., E.M., C.C., T.C., L.S., A.N., A.S.; Methodology: M.R., A.M., and A. R.; project administration: M.R., and A. R.; Resources: A.V., F.C., G.D., P.B., and A.R.; Supervision: A. R., and P. B.; Visualization: M.R., and A.R., Writing – original draft: M.R., S.M., L.S., I.C. and A.R., writing –review & editing: all authors.

## Acknowledgments

This work was supported by MIUR through the PRIN 2017E3A2NR_004 project for A. R., P.B., M.S., S.M., L.S., and F.C. and Center for Colloid and Surface Science (CSGI) through the BOW project, Horizon 2020 – Future and emerging technologies (H2020 – FETOPEN), ID: No. 952183 for P. B. Thanks to PNRR project “Health Extentended ALliance for Innovative Therapies, Advanced Lab-research, and Integrated Approaches of Precision Medicine” - HEAL ITALIA – Spoke 5 – Next Gen Therapeutics - ORGANO-CAR”.

## Data availability statement

Data will be available on request from the authors

## Ethics approval statement

Research was conducted in accordance with The Code of Ethics of the World Medical Association (Declaration of Helsinki) and Good Clinical Practice. The study was approved by the ethics committee of Spedali Civili di Brescia (Protocol number 5705). Patient derived organoids were obtained by Bruno Boerci Oncological Biobank, (ICS Maugeri IRCCS’s ethical committee approvation of 27 July 2009)

## Patient consent statement

Informed consent was obtained from all the subjects enrolled in the study. The privacy rights of subjects involved in the study have always been observed.

