## Supplementary file 1 for "Red Blood Cell-derived Extracellular Vesicles enable Cisplatin and Cetuximab combined Therapy against Triple-Negative Breast Cancer"

<https://orcid.org/0000-0003-2737-1090>

F.Corsi, IRCCS Istituti Clinici Scientifici Salvatore Maugeri, 27100 Pavia, Italy

P.Bergese, National Center for Gene Therapy and Drugs based on RNA Technology, University of Padova, 35122 Padova, Italy

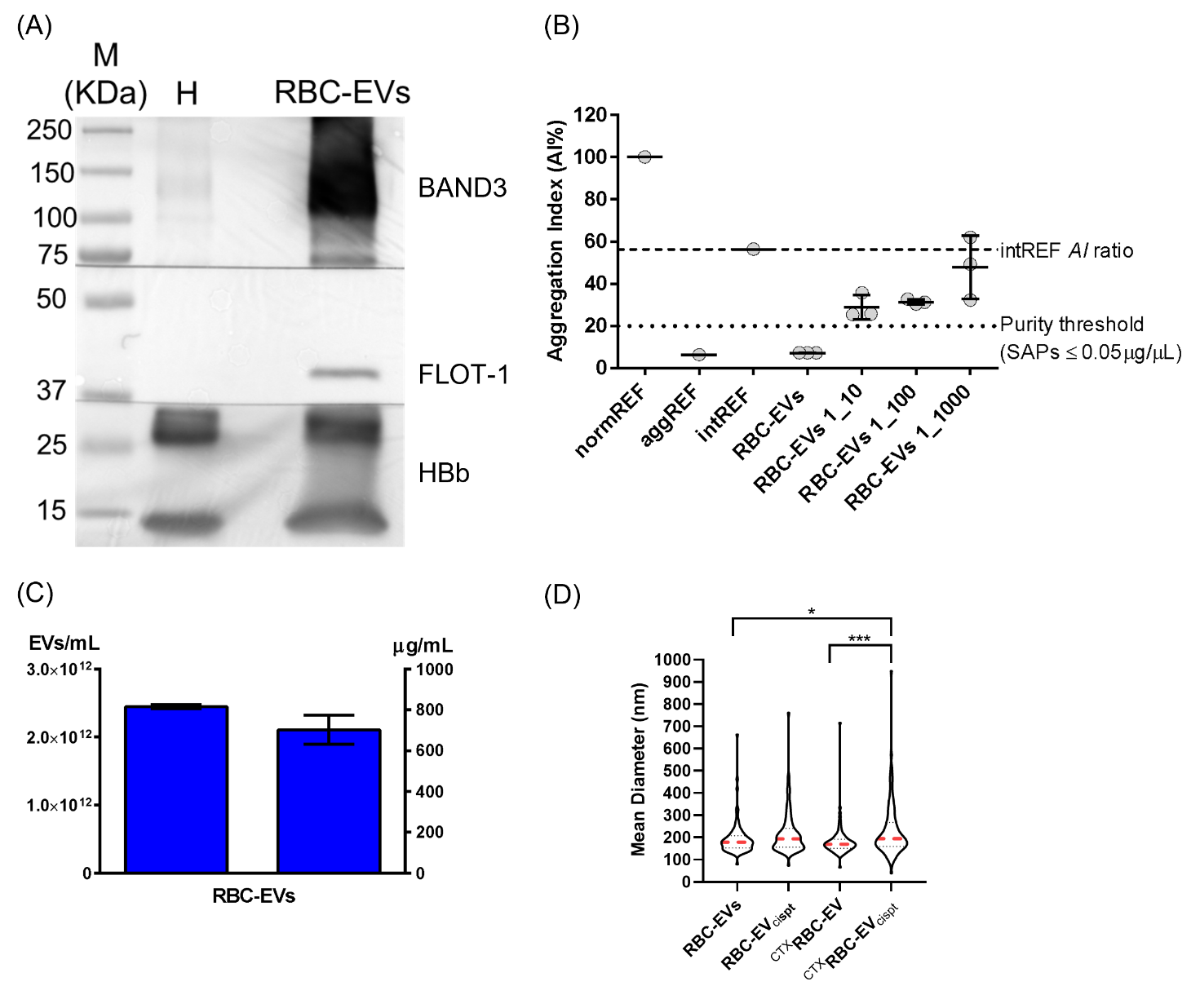

**Figure S1. Biophysical and biochemical characterization of RBC-EVs.** A) Western blot analysis demonstrating the expression of proteins related to RBC-EVs. H, RBC homogenate (30µg); B) CONAN assay on RBC-EV formulations at different dilutions; C) Quantitative analysis of protein and nanoparticle concentration, represented in bar graphs; D) Violin plot obtained with NTA analysis of the different mean size of RBC-EV formulations. One-way Anova: *p = 0.0206, ***p=0.0009.

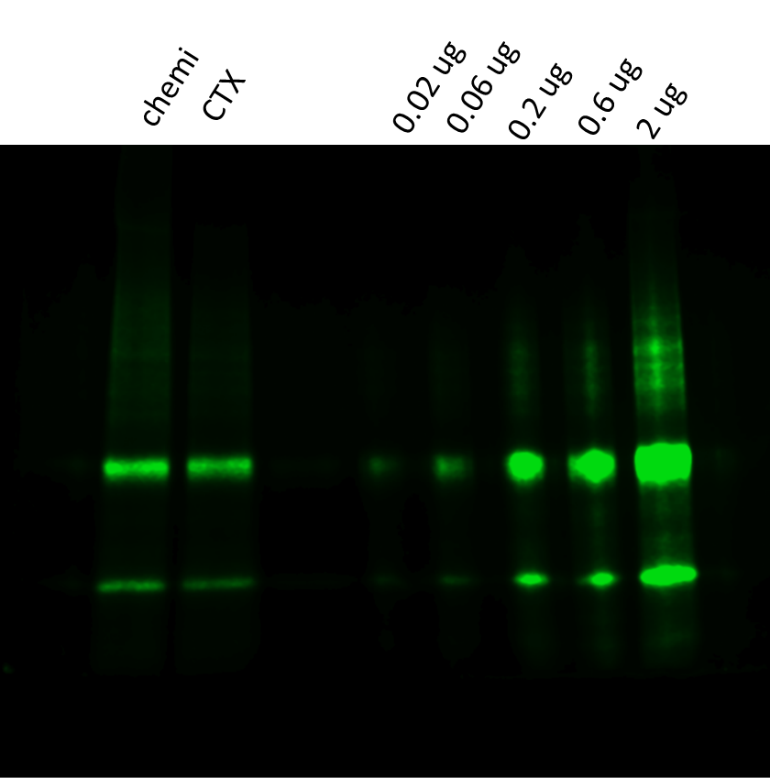

2

0.6

0.2

0.06

0.02

Free CTX (µg)

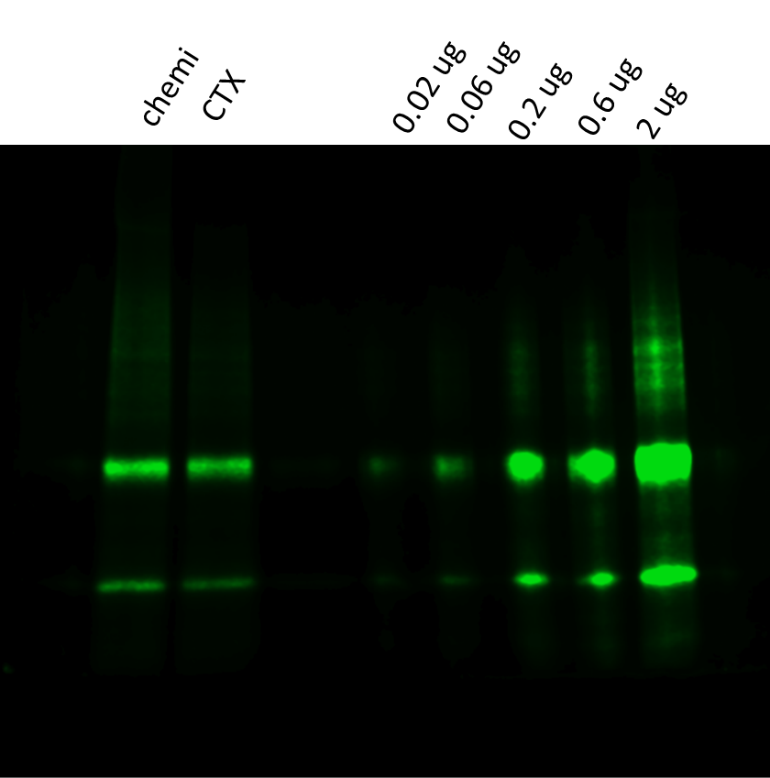

^CTXRBC-EVs^

^CTX High chain^

^CTX Light chain^

**Figure S2. CTX quantification on ^CTX^RBC-EVs.** SDS-PAGE was performed using the Any kD™Mini-PROTEAN TGX Stain-Free precast gel for polyacrylamide gel electrophoresis (24 μL loaded ^CTX^RBC-EVs).

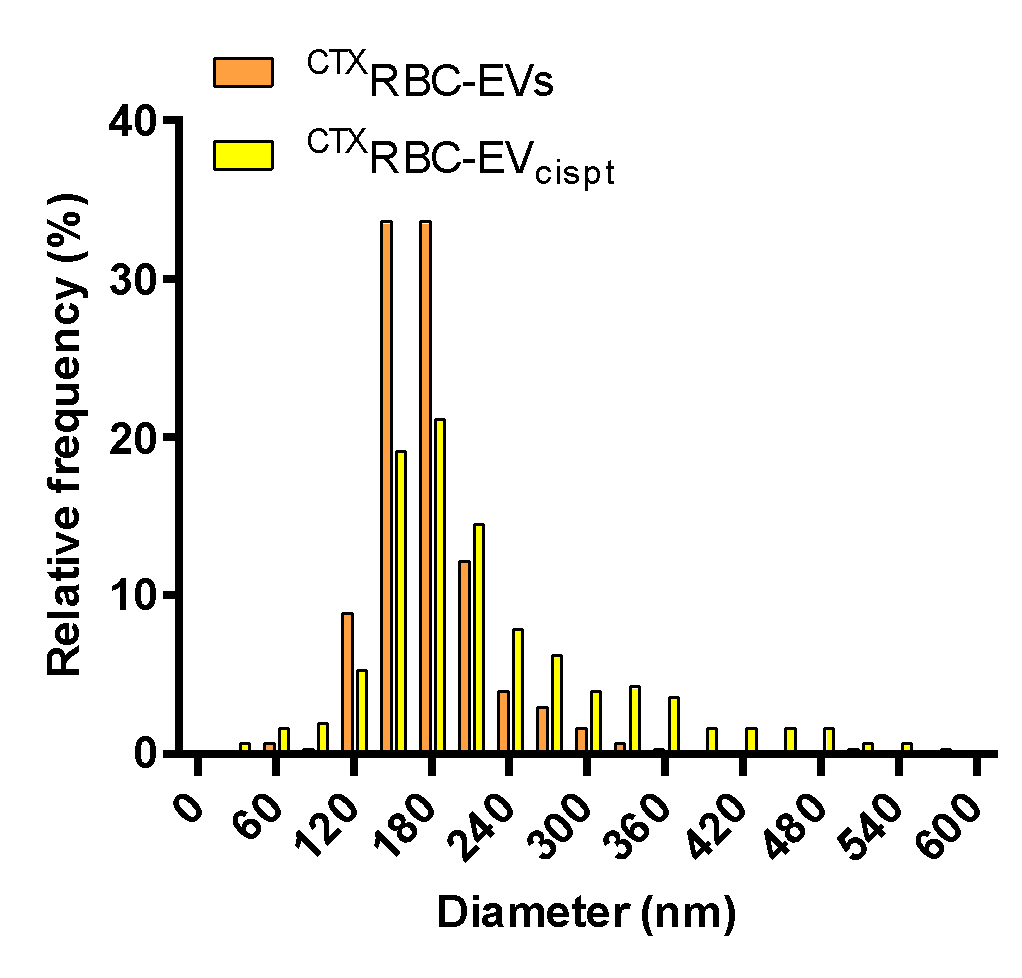

**Figure S3. Comparison between ^CTX^RBC-EVs and ^CTX^RBC-EV_cispt_ size distribution.** Size distributions were measured by NTA, as described in the material and method section.

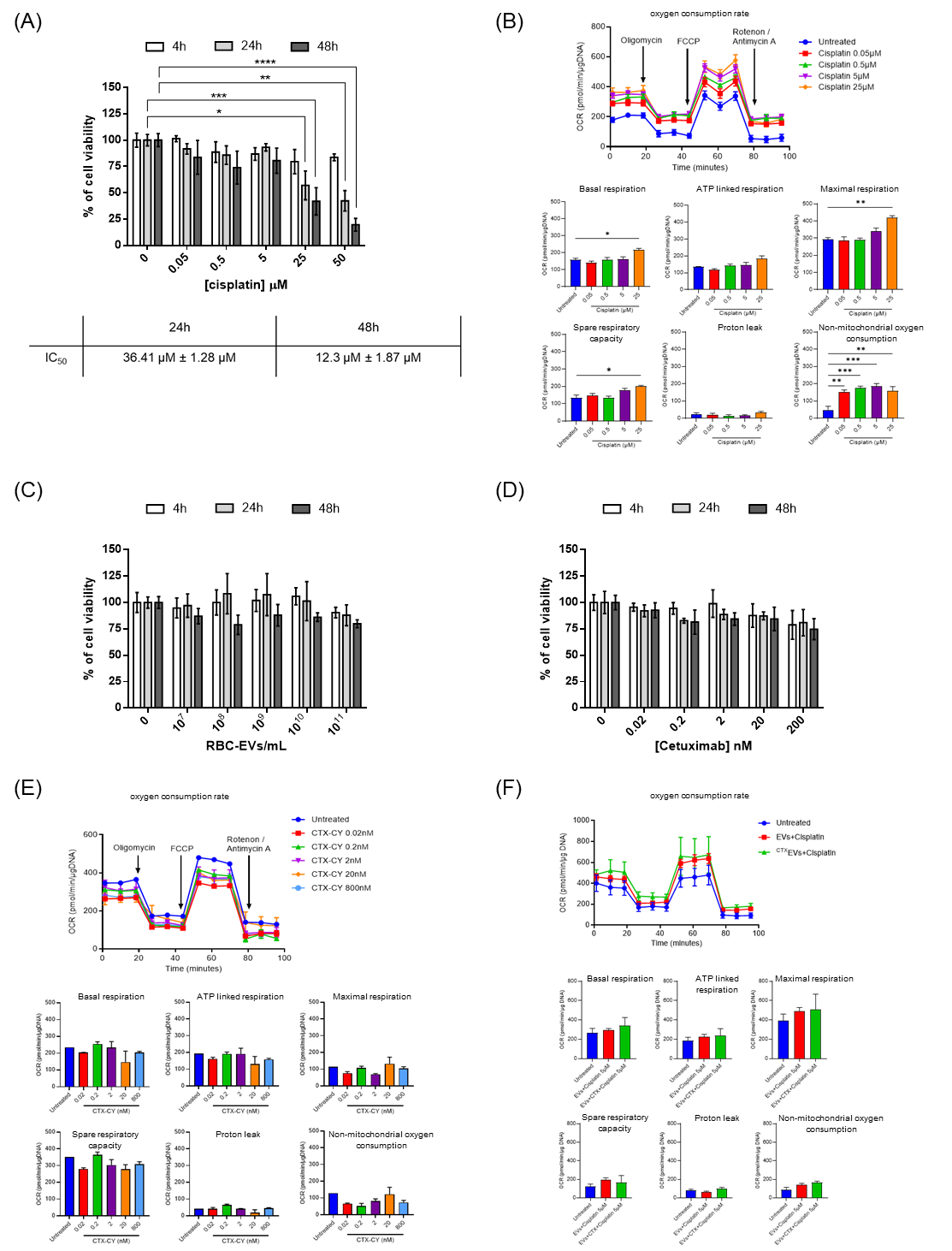

**Figure S4. Assessment of cell viability and metabolic changes of the soluble components.** (A) Cell viability assessment of free cisplatin *via* MTT assay. *p < 0.05, **p ≤ 0.01, ***p ≤0.001, ****p ≤ 0.0001. Data are expressed as mean ± standard error of the mean (SEM). **(B)** Effect of 24h Cisplatin treatment on mitochondrial respiration in MDA-MB-231 cells. OCR values were normalized to DNA content. Data represent mean ± SEM. Statistical analysis was performed with one-way ANOVA and Dunnett’s test comparing each treated group to untreated group, using GraphPad Prism 9.0.0 (San Diego, CA, USA). **(C)** Cell viability assessment of RBC-EVs *via* MTT assay. **(D)** Cell viability assessment of free CTX *via* MTT assay. Data are expressed as mean ± SEM. **(E)** Effect of 24h Cetuximab (CTX) treatment on mitochondrial respiration in MDA-MB-231 cells. OCR values were normalized to DNA content. Data represent mean ± SEM. Statistical analysis was performed with one-way ANOVA and Dunnett’s test comparing each treated group to untreated group, using GraphPad Prism 9.0.0 (San Diego, CA, USA). **(F)** . Effect of 24h treatment with RBC-EV_cispt_ (5µM), and ^CTX^RBC-EV_cispt_ on respiration in MDA-MB-231 cells. Data represent mean ± SEM. Statistical analysis was performed with one-way ANOVA and Dunnett’s test, using GraphPad Prism 9.0.0 (San Diego, CA, USA).

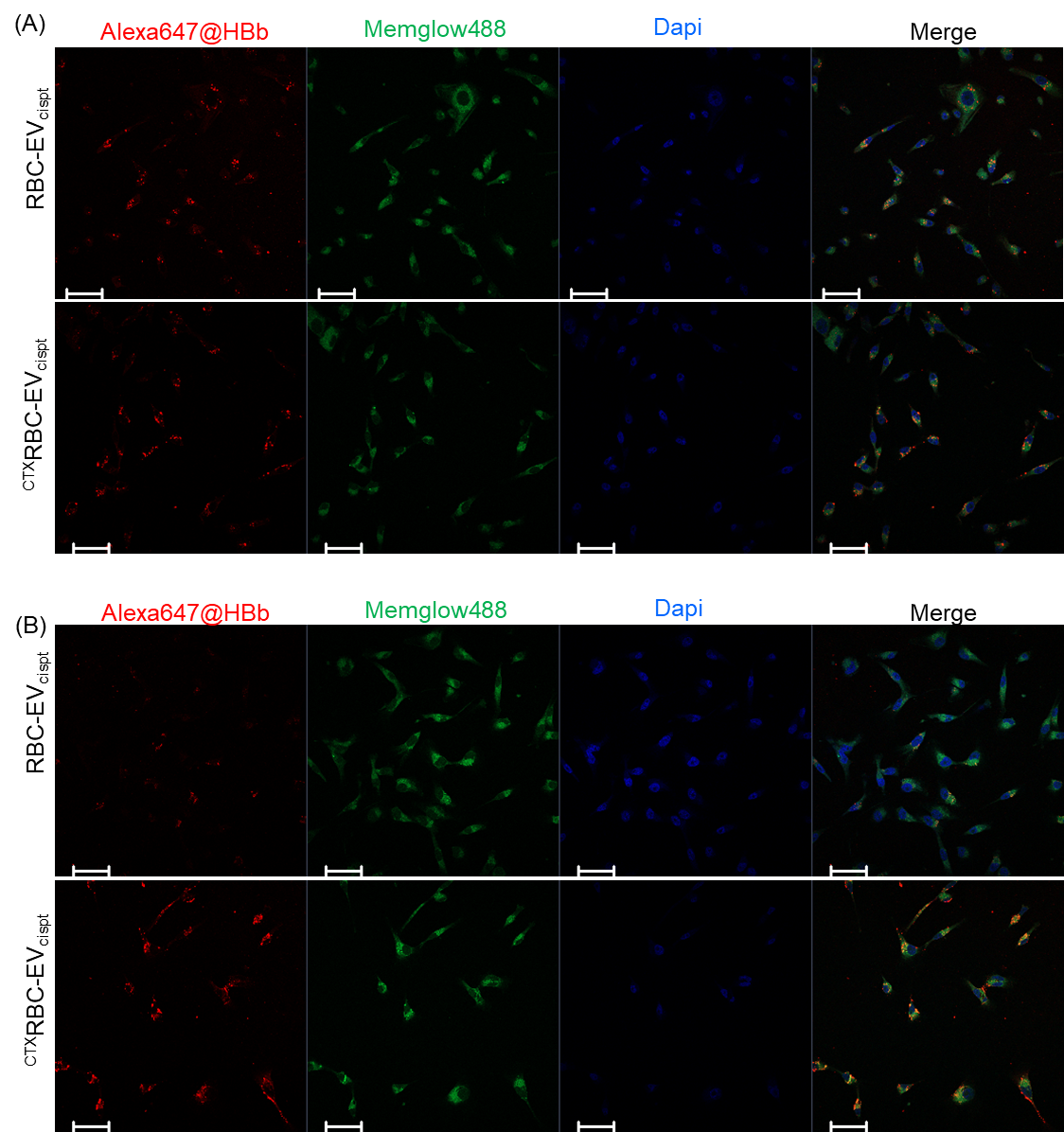

**Figure S5. Uptake of RBC-EV_cispt_ and ^CTX^RBC-EV_cispt_ in 2D TNBC cell model after 4h and 24h.** (A) Uptake after 4h. (B) Uptake after 24h. Scale bar 50µm.

A)

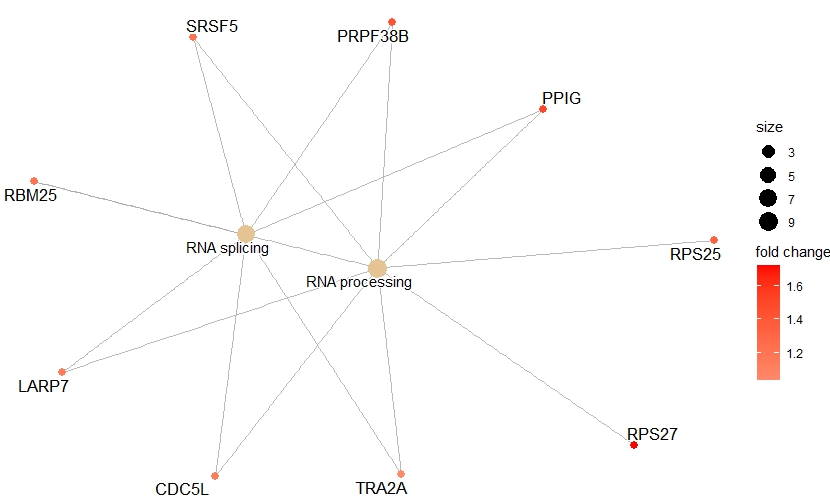

B)

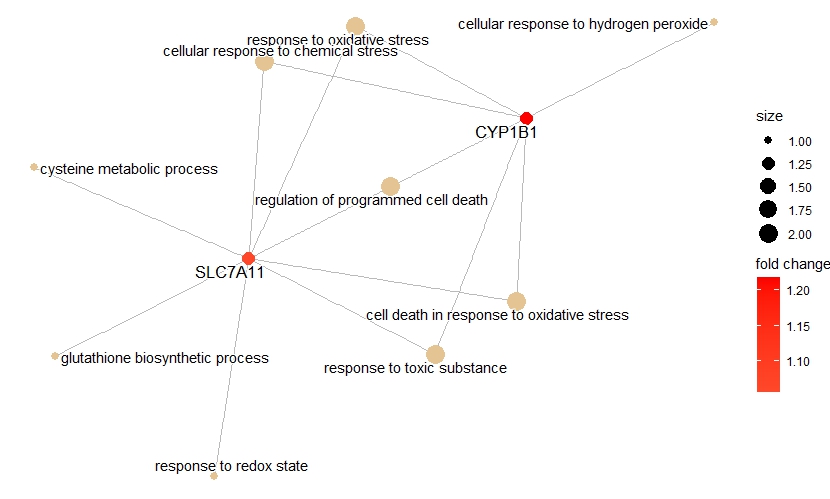

**Figure S6: Biological processes and genes affected by the different treatments.** Network plots show Gene Ontology biological processes (GO-BP) of interest enriched by up-regulated genes in MDA-MB-231 cells treated with (A) RBC-EV_cispt_ and (B) ^CTX^RBC-EV_cispt_. Bubble size of the GO terms represents number of genes involved, while gene color gradient indicates differential expression fold-change (in log2 scale).

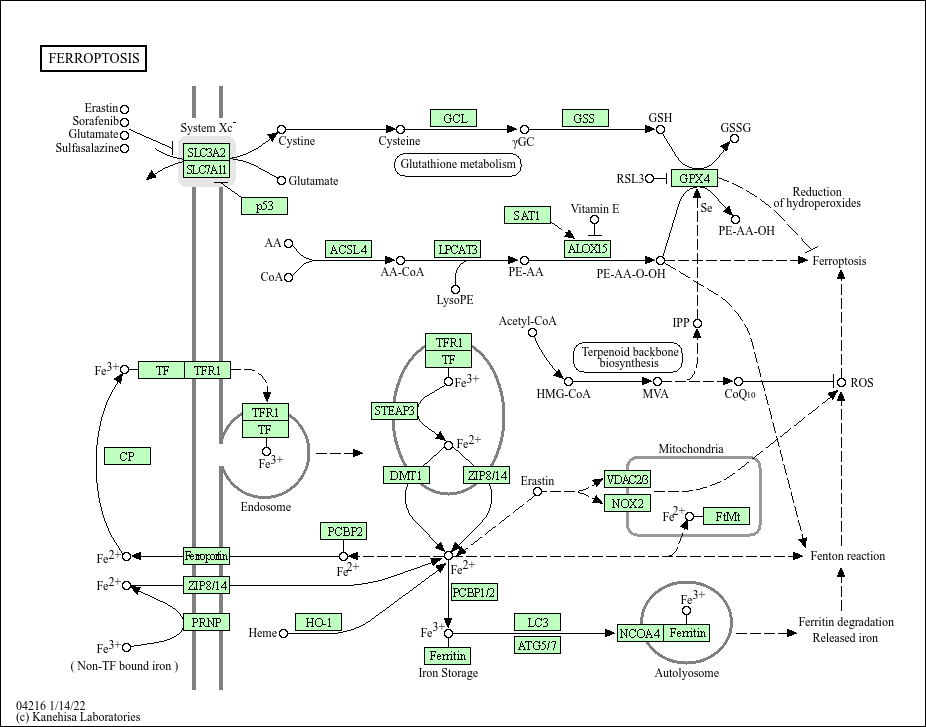

**Figure S7: KEGG map of Ferroptosis Reference pathway (map 04216,**

**https://www.genome.jp/pathway/map04216).**

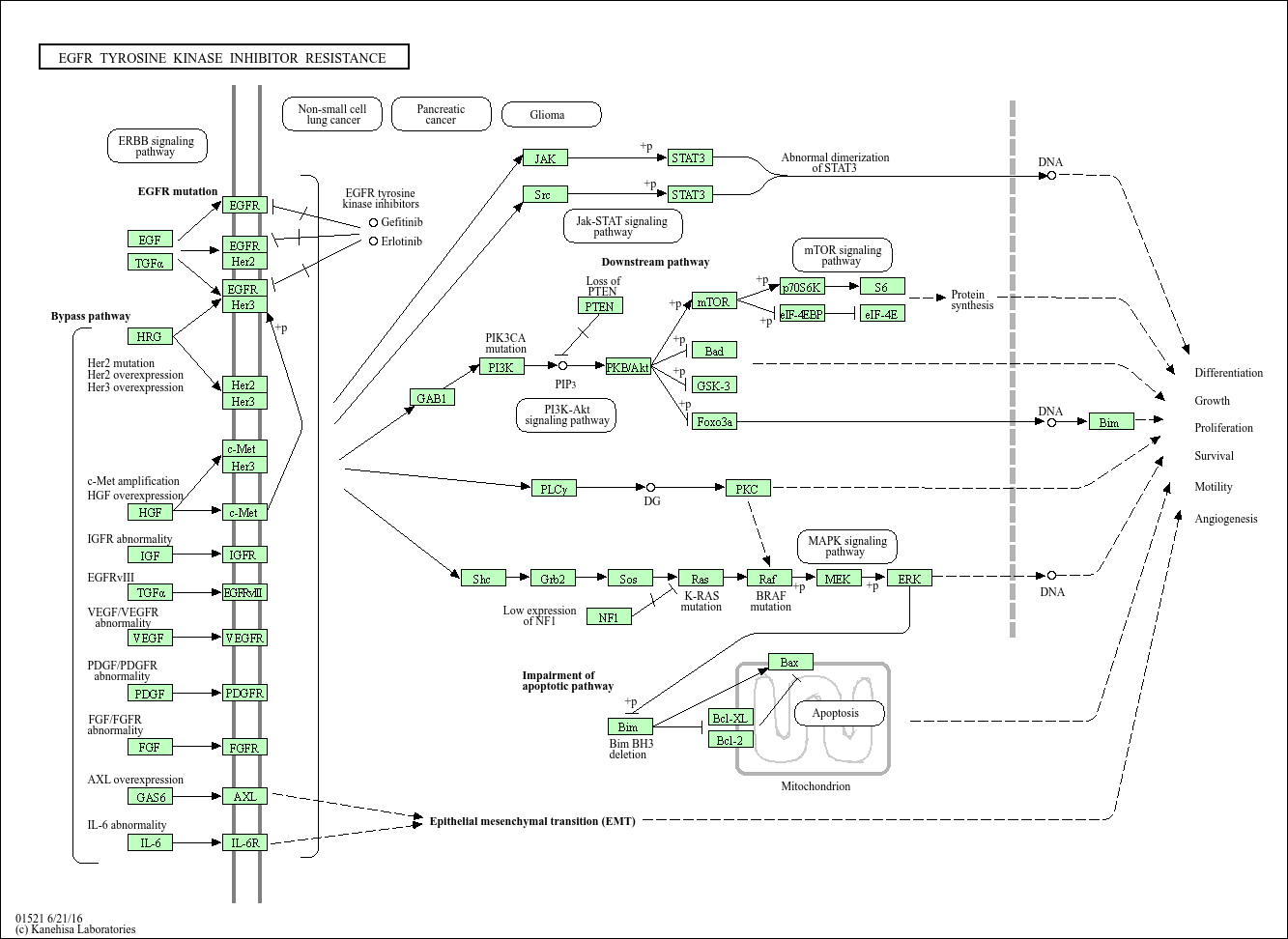

**Figure S8: KEGG map of EGFR tyrosine kinase inhibitor resistance pathway (map 01521,**

**https://www.genome.jp/pathway/hsa01521)**

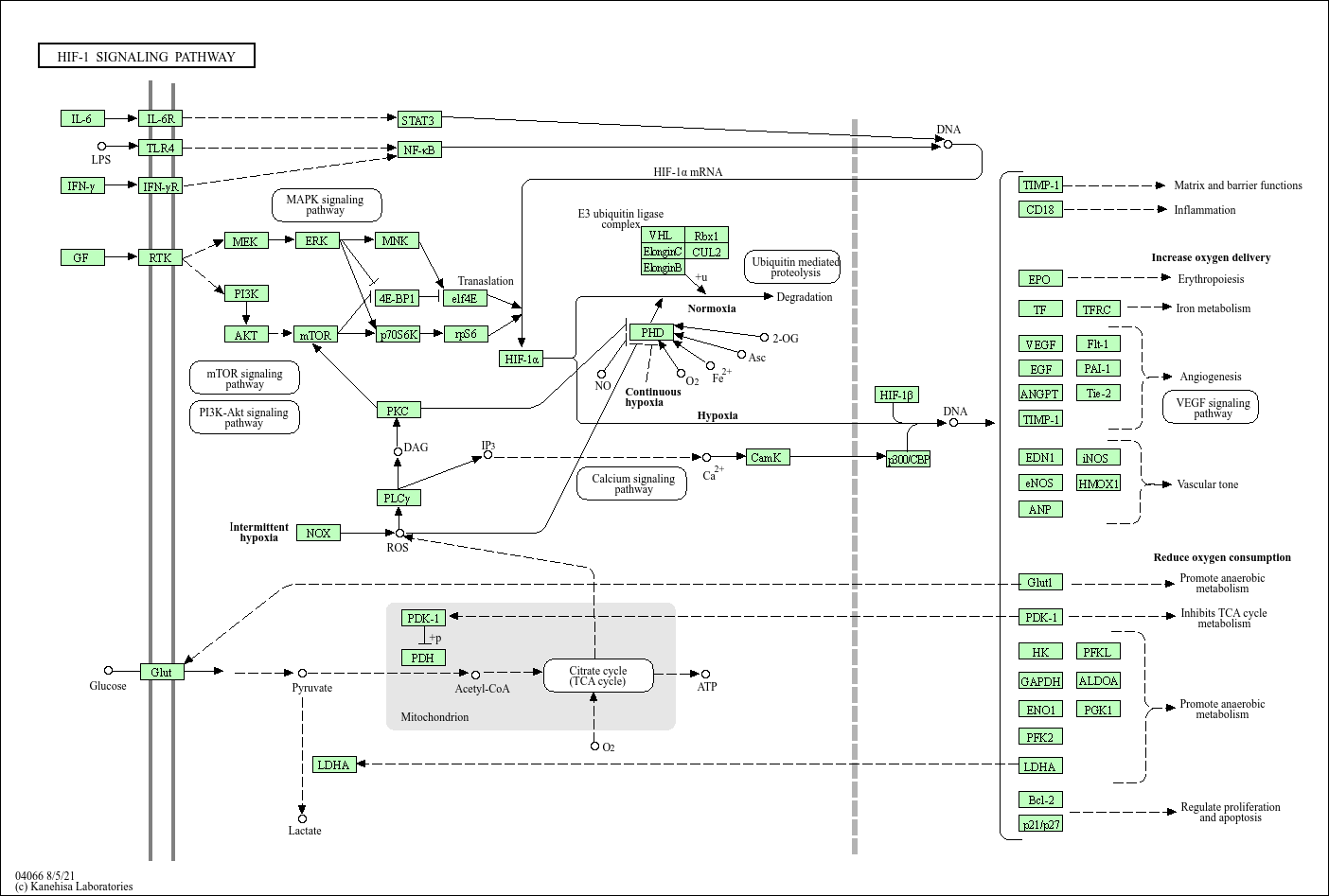

**Figure S9: KEGG map of HIF-1 signaling pathway (map 04066,**

**https://www.genome.jp/pathway/hsa04066)**

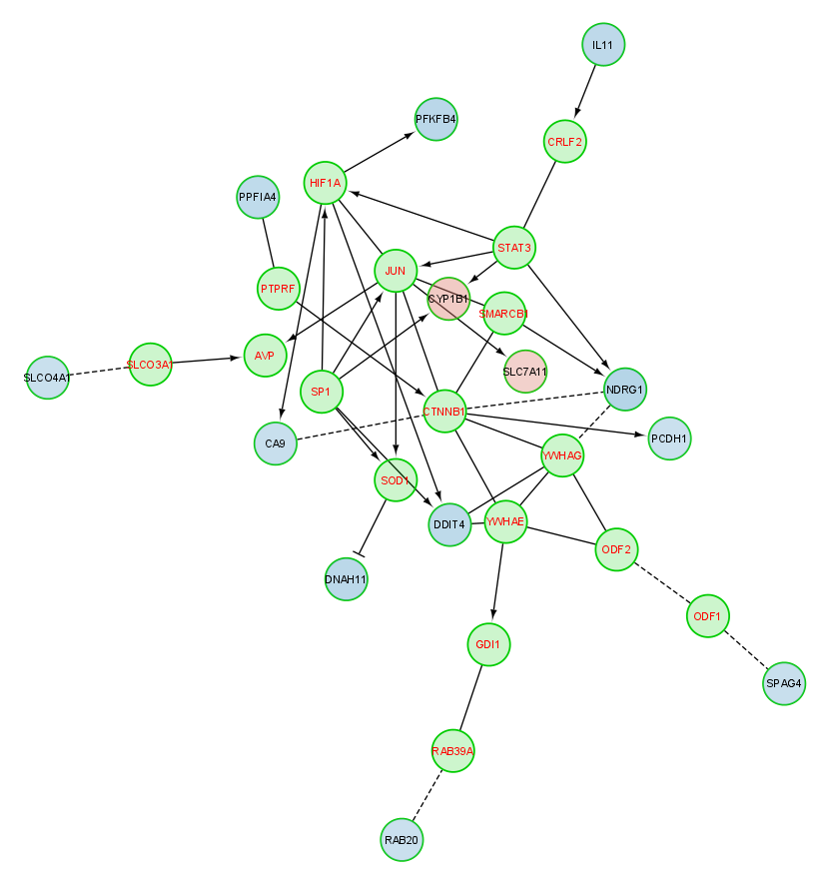

**Figure S10: Cytoscape network of protein-protein interactions among the DEGs found in ^CTX^RBC-EV_cispt_ vs wt treated cells**. Up-regulated and down-regulated genes are represented by red and blue circles, respectively, while green circles indicate linker genes. Network highlights the relationship among STAT3, HIF-1a and its downstream genes.

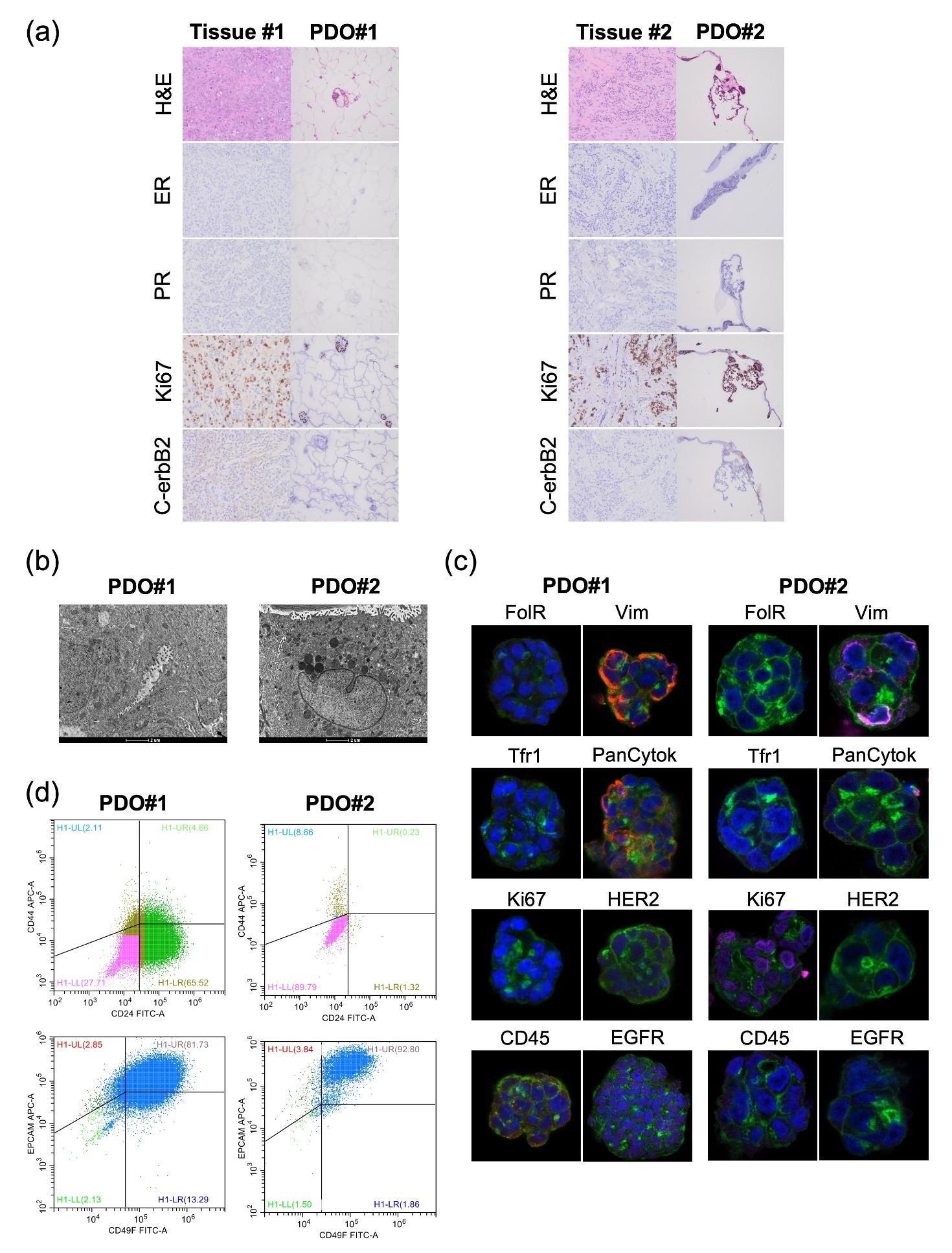

**Figure** **S11. Characterization of the two patient-derived organoids (PDOs), PDO#1 and PDO#2.** (A) Histological and molecular characterization of tumor tissues (Tissue #1 and Tissue #2) and matched PDOs (PDO#1 and PDO#2), by H&E staining and IHC for estrogen (ER) and progesterone (PR) receptors, Ki67 proliferation index and c-ErbB2 receptor. (B) TEM images of PDO#1 and PDO#2, showing tumor-associated morphological features: intracellular lumens rich in microvilli (PDO#1) and mitochondria rich in ridges (PDO#2); scale bar 2 µm. (C) Confocal microscopy analyses assessing the expression of Folate receptor (FolR), Vimentin (Vim), Transferrin receptor 1 (Tfr1), Pancytokeratin (PanCytok), Ki67, HER2, CD45, EGF receptor (EGFR) on both PDO#1 and PDO#2. Nuclei (blue, DAPI), membrane (green, WGA FITC), and markers (pink, anti-Rb AF546) are labelled. (D) PDO#1 and PDO#2 evaluation by flow cytometry for the cell surface markers CD24/CD44 and EpCAM/CD49f.

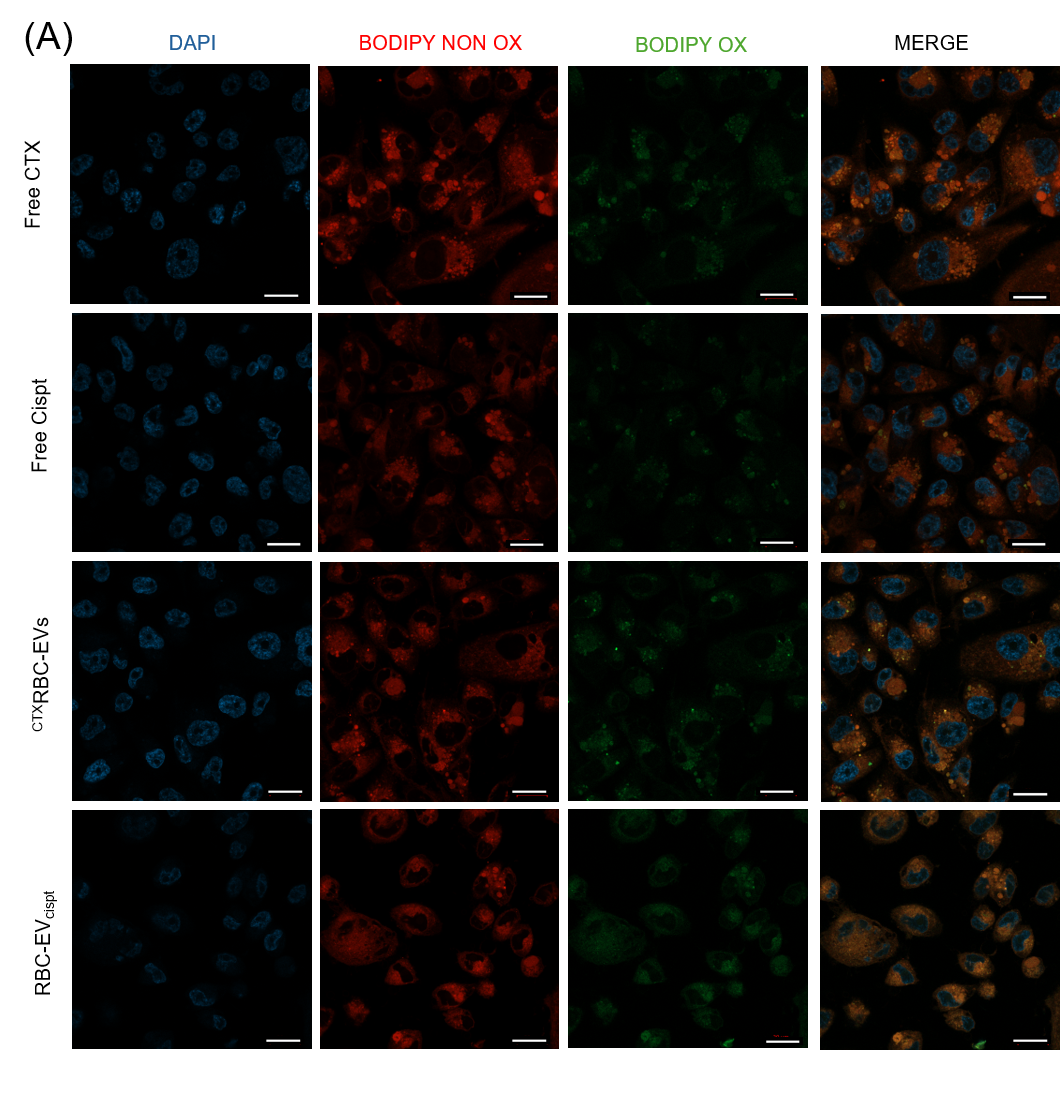

**Figure S12: Ferroptosis promotion evaluation via lipid peroxidation.** (A) Representative confocal microscopy images of cells stained with BODIPY™ 581/591 C11 (oxidized: green; non-oxidized: red) to visualize lipid peroxidation, and DAPI (blue) to label nuclei. Merged images show all channels combined. Treatments include Free CTX, Free Cisplatin (Free Cispt), ^CTX^RBC-EV, RBC-EV_cispt_. Scale bar 20 um.
